# Engineered Peptide Barcodes for In-Depth Analyses of Binding Protein Ensembles

**DOI:** 10.1101/287813

**Authors:** Pascal Egloff, Iwan Zimmermann, Fabian M. Arnold, Cedric A. J. Hutter, Damien Morger, Lennart Opitz, Lucy Poveda, Hans-Anton Keserue, Christian Panse, Bernd Roschitzki, Markus A. Seeger

**Author notes:** These author contributed equally (Pascal Egloff, Iwan Zimmermann).

## Abstract

Binding protein generation relies on laborious screening cascades that process candidate molecules individually. To break with this paradigm, we developed NestLink, a binder selection and identification technology able to biophysically characterize thousands of library members at once without handling individual clones at any stage of the process. NestLink builds on genetically fused barcoding peptides, termed flycodes, which are designed for maximal detectability by mass spectrometry and serve as unique molecular identifiers for accurate deep sequencing. We applied NestLink to overcome current limitations of binder generation. Rare binders against an integral membrane protein were identified directly in the cellular environment of a human pathogen. Hundreds of binder candidates were simultaneously ranked according to kinetic parameters. Adverse effects of target immobilization were overcome by selecting nanobodies against an ABC transporter entirely in solution. NestLink may provide a basis for the selection of tailored binder characteristics directly in tissues or in living organisms.

## INTRODUCTION

Binding proteins have proven invaluable for a plethora of applications in basic science, diagnostics and therapy. Their generation typically involves laborious single-clone experiments, such as ELISA, Sanger sequencing or surface plasmon resonance (SPR), which are needed to identify and characterize individual hits. In recent years, however, novel methods based on next generation sequencing (NGS) and liquid chromatography coupled to tandem mass spectrometry (LC-MS/MS) were introduced, which enabled binder identification directly from ensembles, such as immune repertoires^1–6^. These innovative strategies overcome throughput limitations of single-clone experiments by the following workflow: NGS of a binder pool, capturing of binders at an immobilized target, proteolytic digest of the captured pool members, peptide detection via LC-MS/MS and matchmaking of NGS and LC-MS/MS data for binder identification. Unfortunately, due to extensive sequence homology within binder libraries, the approach is currently limited by the low number of peptides, which are unambiguously assignable to individual binder sequences (most peptides are encoded by several pool members)^7^. Furthermore, many peptides suffer from low ionization and fragmentation efficiencies, thus hampering binder identification significantly.

Here, we present NestLink, a technology that overcomes these inherent limitations and enables for the first time biophysical characterization of thousands of binder candidates without the need to handle individual clones at any stage of the process. NestLink builds on genetically fused barcoding peptides, termed flycodes, which are designed for optimal detectability by LC-MS/MS and which are unambiguously assignable to pool members. Flycodes also serve as unique molecular identifiers (UMIs) for accurate sequence determination by NGS^8, 9^. The dual usage of the flycode sequence at the genetic and the protein level is employed to link genotype and phenotype of library members *in silico*. This enables binder selections from ensembles in analogy to classical display procedures, such as phage display. However, NestLink radically differs from conventional selection methods, as it generates accurate readouts for individual library members. Importantly, NestLink operates in the absence of a *physical* genotype-phenotype linkage and is thus independent of large display particles – a paradigm shift permitting unprecedented, size-dependent selection pressures. In this work, we introduce the basic principle of NestLink and describe three out of many possible applications in binding protein development.

## RESULTS

### The NestLink principle

NestLink centers on a diverse library of short flycodes, which are genetically fused to a library of binding proteins in a novel process termed “library nesting” (Fig. 1). The nested library is sequenced by NGS to assign all flycodes to their corresponding binders (Fig. 1, Supplementary Fig. 1). Subsequently, the nested library is expressed as a pool and subjected to selection pressures in the absence of the genotype. Flycodes of selected binders are isolated via sequence-specific proteases and detected via LC-MS/MS (Supplementary Fig. 2). In combination, library nesting, NGS and LC-MS/MS establish an *in silico* genotype-phenotype linkage, which allows rapid characterization of individual binder properties.

**Figure 1:**
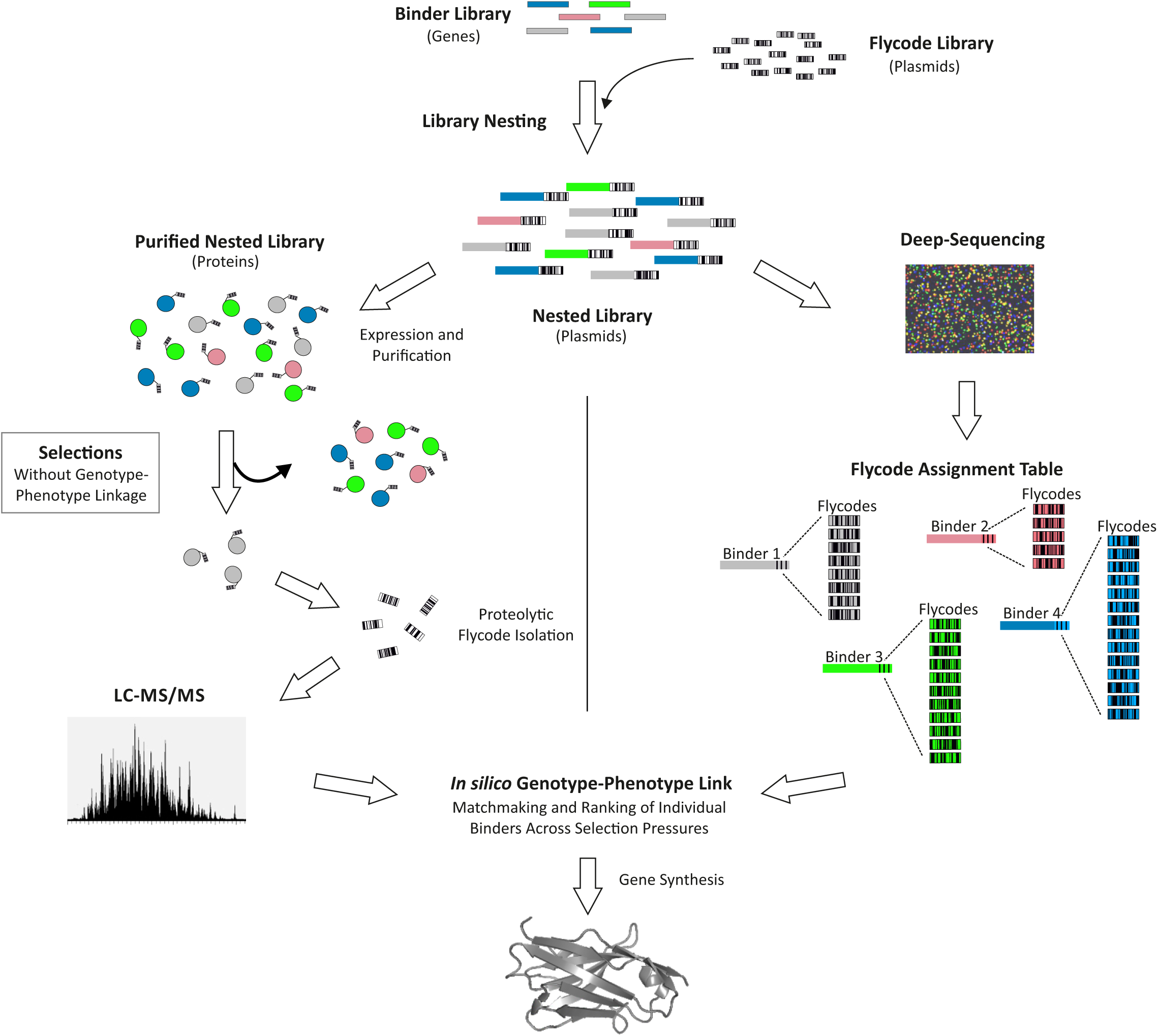
NestLink overview. Details are provided in the main text.

### Library nesting

Library nesting links each gene of a binder library in a controlled manner multiple times to unique flycodes (Fig. 2). In a first step, a defined number of bacterial colony forming units (cfu), harboring plasmids that encode binder library members, are pooled for plasmid isolation. This step defines the maximal diversity of the binder library under investigation. In a second step, restriction digest and ligation is used to clone the binder library into a plasmid, which harbors the flycode library. Thereby, the binder library and the flycode library are nested. The number of cfu pooled for plasmid isolation in this second step defines the maximal number of flycodes under investigation and thus the average number of flycodes per binder. For example, if the binder library size is < 1,000 and 30,000 cfu are pooled after the library nesting step, the number of different flycodes per binder is on average > 30. Importantly, attached flycodes are unique, because the experimental flycode library diversity (≈ 100 Mio) vastly exceeds the total number of flycodes linked via library nesting. Hence, flycodes are unambiguously assignable to library members, which is the basis for unambiguous binder detection via LC-MS/MS. Of note, library nesting avoids PCR amplification and thereby prevents undesired recombination events (Supplementary Fig. 3).

**Figure 2:**
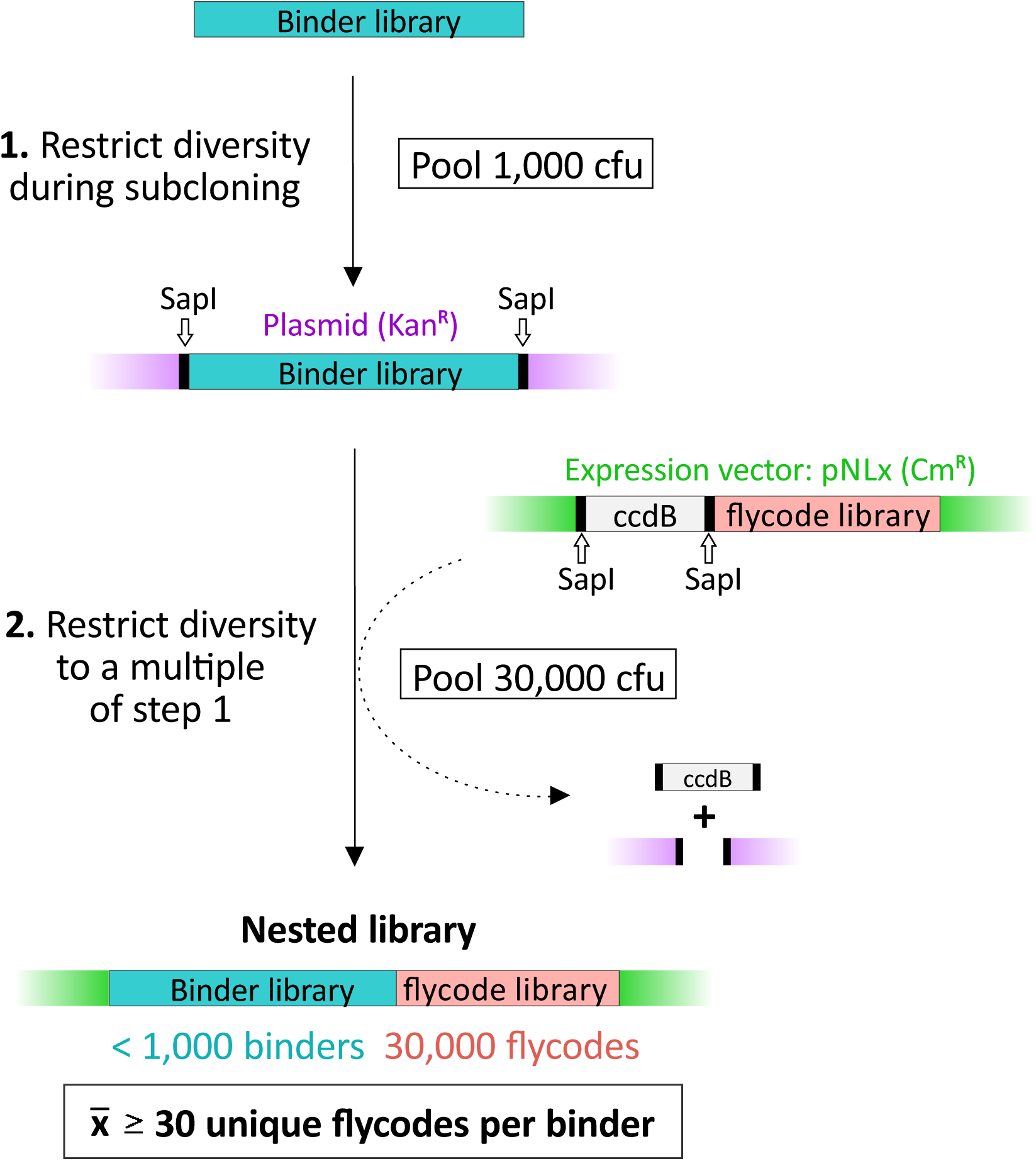
Library nesting. This example shows how a nested library containing on average 30 flycodes per binder is generated. First, the maximal binder diversity is restricted to 1,000 by isolating 1,000 cfu of bacteria harboring the binder library on a plasmid. This restricted binder pool is cloned into a vector backbone encoding for the flycode library. By isolating 30’000 cfu of the resulting nested library, the maximal number of flycodes is restricted to 30’000 and each binder on average is fused to 30 different flycodes. The large diversity of the flycode library (100 Mio) ensures uniqueness of attached flycodes (30,000 << 100 Mio).

### Flycode library design

The flycode library is composed of genetically encoded peptide sequences designed for optimal detection via LC-MS/MS upon proteolytic isolation from a protein pool of interest (Fig. 3a). Flycodes are 11-15 amino acids long and contain two randomized regions resulting in a theoretical library diversity of 5.3 x 10^8^. To enable optimal detection of individual flycodes by LC-MS/MS, the library was designed to be maximally diverse in terms of (i) mass-over-charge ratios (m/z) to fall into the optimal m/z-detection window of high-field orbitraps (550-850 m/z), and (ii) hydrophobicity, thus exploiting the full separation capacity of a typical reverse-phase liquid chromatography system (Fig. 3b, Supplementary Fig. 4a). Flycodes contain an invariant arginine as sole positively charged residue, which supports efficient ionization. The randomized regions are devoid of cysteines and methionines to avoid oxidation and cross-linking, but frequently contain aspartate and glutamate to enhance solubility, as well as proline to facilitate collision-induced fragmentation. Importantly, size exclusion chromatography (SEC) analyses revealed that the attachment of flycodes does not change the oligomeric state of binders (Supplementary Fig. 5). As individual library members are fused redundantly to multiple flycodes, potential negative effects of individual flycodes are averaged out.

**Figure 3:**
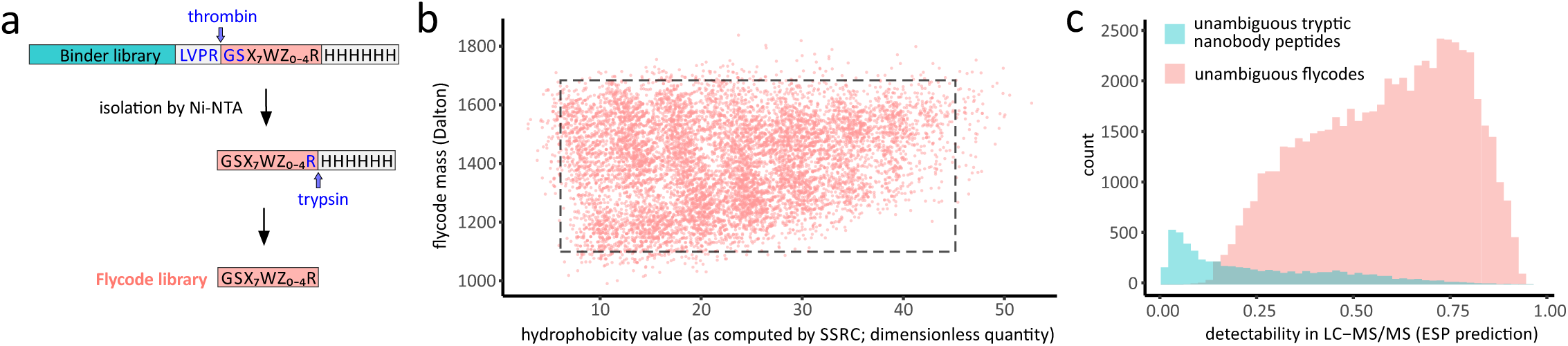
Flycode library design and characteristics. (**a**) Flycodes are located between a thrombin cleavage site (blue) and a His-tag, which can be cleaved site-specifically by trypsin at the sole positively charged amino acid (blue arginine). “X7” denotes a stretch of seven randomized amino acids, and “Z0-4” represents five distinct sequences of 0-4 amino acids in length. Amino acid compositions are provided in the online methods. (**b**) Prediction of hydrophobicity (SSRC; Sequence Specific Retention Calculator14) and parent ion mass for 10,000 randomly chosen flycodes. The optimal detection window for the exclusively doubly-charged flycodes is shown as dashed rectangle. (**c**) Histogram showing the detectability of unambiguously assignable tryptic nanobody peptides (cyan) and unambiguously assignable flycodes (pink) from the same nested library consisting of 3,390 unique nanobodies linked to 59,974 flycodes (see application II, Fig. 5). Peptides are binned according to their ESP prediction value (high ESP values correlate with better detection by LC-MS/MS).

In order to assess the benefits of flycode-mediated protein detection by LC-MS/MS, a nested library comprising 3,390 unique nanobodies linked to 59,974 flycodes (see also application II below) was analyzed *in silico* to compare the flycodes with the peptides obtained by tryptic digest of the nanobodies (Fig. 3c). The analysis revealed 10 times more flycodes than tryptic nanobody peptides that are unambiguously assignable to a single nanobody of the library. In addition, the enhanced signature peptide (ESP) predictor indicates an overall high MS/MS-detectability of flycodes, while a large fraction of the tryptic nanobody peptides are predicted to be poorly detectable^10^. Titration experiments using defined flycode sets revealed that individual binder concentrations can be proportional to the summed MS1 intensities of the corresponding flycodes over more than two orders of magnitude (Supplementary Fig. 6).

### Application I: ranking hundreds of binders according to their off-rates

The identification and biophysical characterization of binding proteins is the most laborious step of the binder generation cascade, as it requires the analysis of individual binder candidates. We applied NestLink to simultaneously characterize more than a thousand synthetic nanobodies (sybodies), which were previously enriched by *in vitro* display against maltose-binding protein (MBP) (Zimmermann et al., under review). To this end, a pool of around 1,200 sybodies was nested with approximately 12,000 flycodes, as determined by estimating cfu numbers. The flycodes were assigned to the respective sybodies using overlapping, paired-end Illumina sequencing with 2×300 bp-reads. NGS covered the complete nested library (442-454 bp) at an average raw-read redundancy of 54 per flycode and 682 per sybody. In addition to standard filtering criteria, the flycode sequences were used as UMIs, i.e. binder sequences linked to the same flycode were aligned and only sequences exceeding a stringent consensus score threshold were considered further (see online methods)^8^. This corrected for most sequencing errors and excluded rare ambiguous flycodes, which were attached to more than one sybody (Supplementary Fig. 1). As intended by library nesting, NGS revealed 12,160 unique flycodes linked to 1,070 unique sybodies (on average 11.4 flycodes per binder).

The nested library was expressed as an ensemble in *E. coli*, purified via His-tag and the monomeric binder candidates were isolated by SEC (Fig. 4a). Thereby, the binder pool was exposed to selection pressures that are impossible to apply in the presence of large display particles, used for example in phage-, ribosome- or yeast display. The monomeric nested library members were mixed with MBP, followed by a second SEC run to isolate MBP-sybody complexes. The complexes were immobilized via biotin previously attached to MBP on two streptavidin-sepharose spin-columns with the aim to identify sybodies exhibiting slow off-rates. To this end, we washed one spin-column with buffer containing a large excess of non-biotinylated MBP, thereby removing binders with fast off-rates, while the other column was washed with buffer only. Flycodes of sybodies that remained on the spin-columns were isolated and analyzed by LC-MS/MS (one LC-MS/MS run per spin-column). The flycode MS1 intensities were assigned to the respective sybodies using the flycode assignment table obtained from NGS. Of 1,070 nested sybodies, the majority (IDs: 0-872) was not detected on either spin-column, presumably because they were either poorly expressed, not monomeric or not forming a stable complex with MBP (or a combination thereof). Eighty-six sybodies (IDs: 873-958) were only detected on the unchallenged spin-column, suggesting fast off-rates. 112 sybodies (IDs: 959-1,070) were detected on both columns and their summed MS1 intensities were used to determine for each binder the fraction remaining on the challenged column compared to the control column (Fig. 4b). To validate the NestLink result, we synthesized several genes of selected sybodies, expressed and purified them individually and determined their off-rates by SPR (Supplementary Fig. 7). In agreement with the applied selection pressures, all tested sybodies were well expressed and monomeric when purified individually. Importantly, off-rates strongly correlated with the fraction remaining on column as determined by NestLink (Fig. 4c). Thus, NestLink allowed us to directly rank well-expressed and monomeric candidates according to their off-rates.

**Figure 4:**
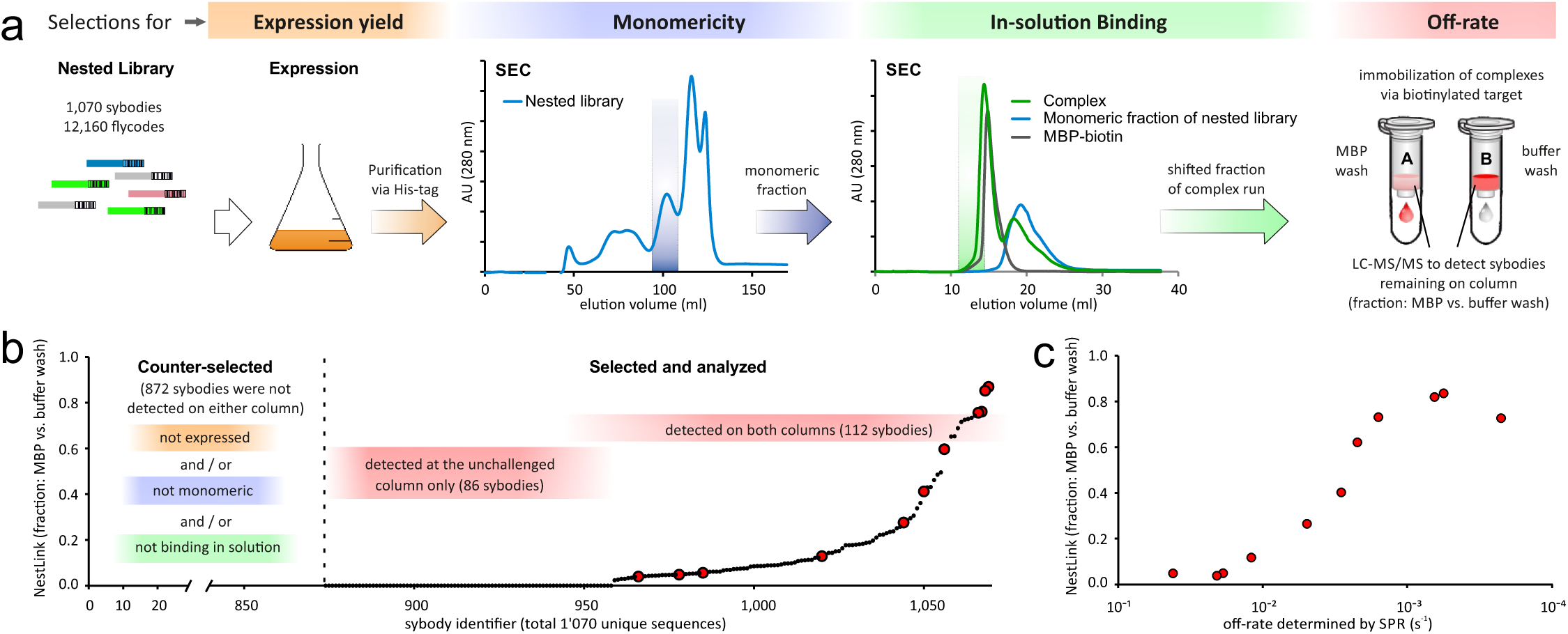
Application I: ranking hundreds of binders according to their off-rates. (**a**) A pool of 1,070 synthetic nanobodies (sybodies) previously enriched against maltose binding protein (MBP) was linked to 12,160 flycodes, as determined by NGS. The nested library was expressed in *E. coli* (orange), purified and separated by SEC (blue). The monomeric pool members were mixed with MBP-biotin and the binders co-migrating with the target on SEC were immobilized on two streptavidin-sepharose columns (red). An off-rate selection was performed by washing one column with buffer containing an excess of non-biotinylated target (MBP wash), while the other column was not challenged (buffer wash). Flycodes linked to sybodies that remained on the columns were isolated and analyzed by LC-MS/MS (one run per column). (**b**) Individual sybodies ranked according to their relative fraction remaining on the MBP-washed column versus the unchallenged column, as determined by the sum of flycode MS1 intensities for each identified binder. (**c**) Individual sybody genes (red, enlarged data points in (b)) were synthesized, followed by expression, purification and SPR-characterization. The recorded off-rates (x-axis) strongly correlate with the fractions remaining on the columns as determined by NestLink (y-axis).

### Application II: nanobody selections without target immobilization

All currently available binder generation strategies rely on target immobilization, which can lead to inaccessibility of epitopes and the enrichment of binders that interact non-specifically with surfaces. We suspected problems of this kind during our previous efforts to generate crystallization chaperones for the bacterial ABC transporter TM287/288^11, 12^. In brief, we immunized an alpaca with detergent-purified transporter and performed two rounds of phage display, ELISA screening and Sanger sequencing of hits^13^. Although we obtained and sequenced 210 specific ELISA hits, only 33 unique, often nearly identical nanobody sequences belonging to merely 5 binder families were identified, which cover a limited epitope space.

To overcome potential selection biases associated with target immobilization, we replaced phage display, ELISA and Sanger sequencing by NestLink, which permits selections entirely in solution via SEC, due to the absence of large display particles. To this end, we linked 3,390 unique nanobody sequences from B-cells of the same immunized alpaca to 59,974 flycodes by library nesting (Fig. 5a). NGS revealed that the nanobodies were present at very different levels in the repertoire, as manifested by a wide range of flycode numbers linked to the 3,390 unique nanobody sequences (Supplementary Fig. 4b). The nested library was purified and all monomeric library members were subsequently mixed with TM287/288 at three different ratios before separation by SEC. Hereby, three distinct levels of pool-internal competition for target binding in solution were achieved, representing unprecedented selection pressures unamenable to conventional display procedures (Fig. 5b). Flycodes were isolated from the nanobody-TM287/288 complex fractions of the three SEC runs and from a sample of the purified nested library (selection input). Subsequently, they were analyzed in independent LC-MS/MS runs. The analysis revealed a large number of efficient binders, which gained in relative abundance at the target as a consequence of increasing pool-internal competition. In total, we identified 29 binder families – more than 5-fold the number of families obtained by the conventional workflow using phage display, ELISA and Sanger sequencing (Fig. 5c). The NestLink outcome was validated by SPR for 11 individually synthesized and purified nanobodies belonging to 11 different families. Specific binding with affinities down to the picomolar range was observed for 9 nanobodies (Fig. 5d, Supplementary Fig. 8). Two binders did not exhibit target binding in SPR. Of note, the two binders belong to families that were also identified via the conventional phage display approach, and in spite of strong ELISA signals, target interaction in SPR was not detectable for any member of these families. The successful binder identification via two orthogonal approaches suggests that the missing signal in SPR is a false-negative result. In agreement with NestLink, four control nanobodies that were well detected in the purified input pool, but not at the target, exhibited no binding in SPR. In summary, these results show that NestLink enables efficient binder identification directly from immunized animals without potential biases resulting from large display particles and/or target immobilization. Furthermore, we show that the technology can even detect binders present at extremely low frequencies in the input pool (0.017 – 0.075 %), suggesting a superior diversity mining capacity for NestLink over current state-of-the-art methods.

**Figure 5:**
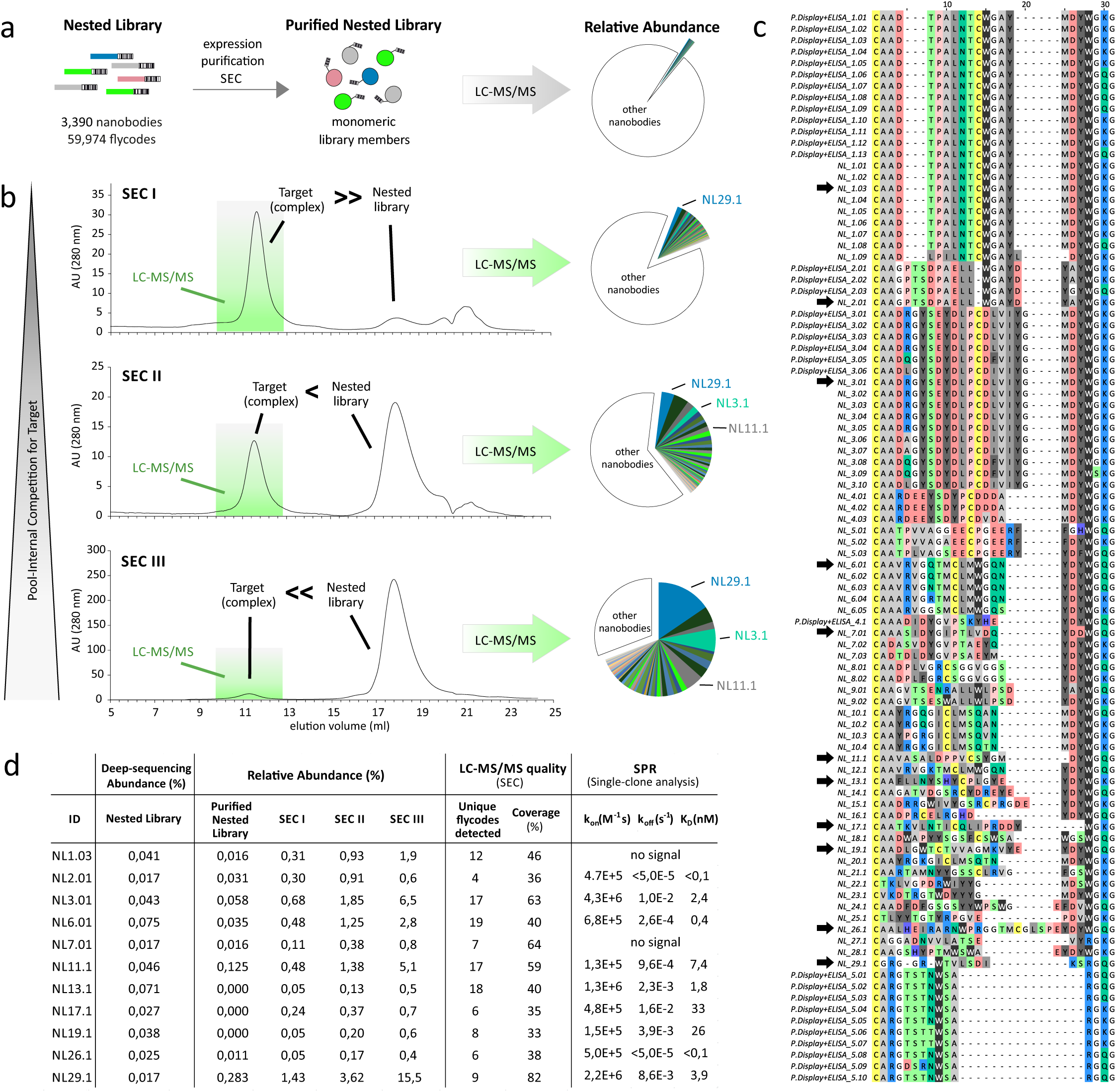
Application II: nanobody selections without target immobilization. (**a**) 3,390 unique nanobody sequences from B cells of an alpaca that was previously immunized with the ABC transporter TM287/288, were nested with 59,974 flycodes, followed by expression, purification and SEC separation of the nested pool to isolate monomeric binder candidates. (**b**) Pool-internal competition for target binding in solution was applied by mixing the nested nanobody pool and TM287/288 at different ratios prior to complex isolation by SEC separation (SEC I – III). Four separate LC-MS/MS runs were performed to analyze the flycodes of the purified nested library as well as of the target bound binders of the three SEC runs. The relative abundance of individual nanobodies within the same LC-MS/MS run was determined according to the summed MS1 intensities of the detected flycodes. The colored slices represent 61 individual nanobodies that gained in relative abundance as a consequence of increased competition for the target. Other nanobodies that did not gain in abundance are collectively represented by the colorless slices. (**c**) CDR3 alignment of the 61 unique nanobodies identified via NestLink (NL1.01 – NL29.01) or via phage display and extensive ELISA screening using the same immunized animal (P. Display + ELISA_1.01 – 5.10). Black arrows denote clones that were characterized individually by SPR. (**d**) Comparison of NestLink and SPR results.

### Application III: specific recognition of an outer membrane protein in the cellular context

*In vitro* selections against detergent-purified integral membrane proteins can generate a large number of high-affinity binders. However, they often fail to bind in the cellular context, due to inaccessibility of epitopes or due to different conformational states adopted by the target in the lipid environment. To provide a solution to this common roadblock, we applied NestLink for deep-mining a pool of sybodies, which was previously enriched by *in vitro* selections against the detergent-purified major outer membrane protein (MOMP) of the human pathogen *Legionella pneumophila serogroup 6* (*Lp-SG6*). ELISA revealed that 12 % of the display-derived pool exhibited specific interactions to the target in detergent. However, flow cytometry experiments using atto488-labelled sybodies suggested that none of the hits recognized the target in the cellular context of *Lp-SG6*, where MOMP is embedded in a dense layer of lipopolysaccharides (LPS). This indicated that desired binders recognizing MOMP in the cellular context were heavily underrepresented or entirely absent.

For pool-mining, we generated a nested library encoding 1,444 unique sybodies linked to 23,598 unique flycodes, as determined by NGS. Subsequently, we selected monomeric sybodies by SEC and performed a pull-down with intact *Lp-SG6* or with one of the three control strains *Escherichia coli*, *Citrobacter freundii* or *Lp-SG3* (Fig. 6a). Captured sybodies were subsequently analyzed via their corresponding flycodes by independent LC-MS/MS runs, which allowed monitoring of relative binder abundances on the four bacterial strains. From the initial 1,444 sybodies, 157 passed the pre-selection for monomericity *and* were unambiguously detected at one or several of the four bacterial strains. Interestingly, only five rare sybodies (representing between 0.05-0.22 % of the pool) exhibited a pronounced increase in relative abundance at *Lp-SG6,* as compared to the input material of the pull-down, whereas for the other three bacterial strains no significant increase for any of the 157 sybodies was observed (Fig. 6b). As *Lp-SG6* was the only cell type of the pull-down that harbored the MOMP-variant used for the initial *in vitro* selection, this result confirmed the excellent specificity and sensitivity of NestLink (Fig. 6c).

**Figure 6:**
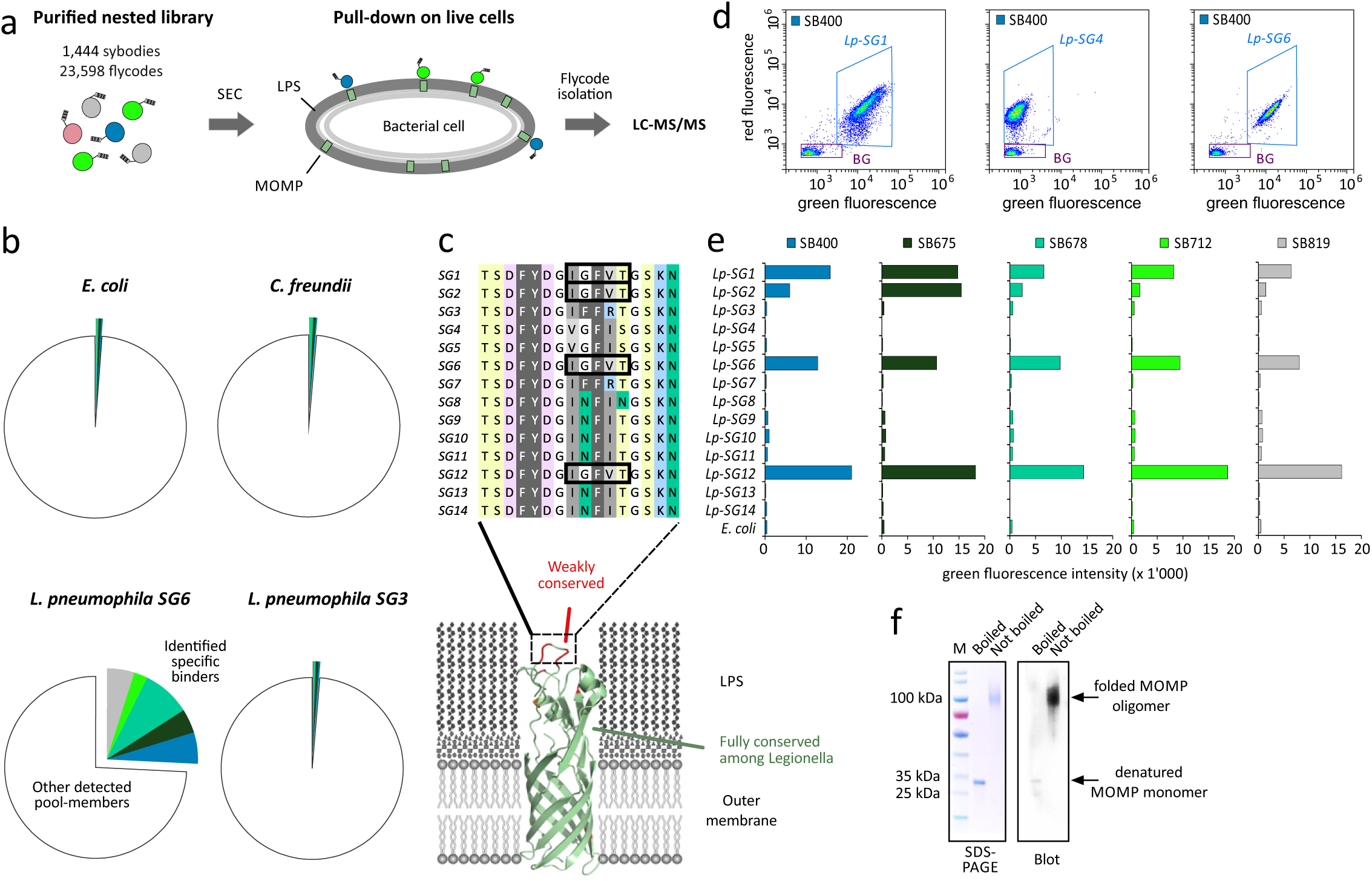
Application III: specific recognition of an outer membrane protein in the cellular context. (**a**) A nested library was constructed from a sybody pool that was previously enriched by *in vitro* selections against detergent-purified MOMP from *Lp-SG6*. After isolating monomeric nested library members by SEC, sybodies recognizing MOMP embedded in the outer membrane of *Lp-SG6* were selected by a pull-down on intact cells. (**b**) LCMS/MS was used to monitor the relative abundance of each sybody on the target cell, as well as on three control strains (*E. coli, C. freundii, Lp-SG3)*. Five specific sybodies exhibiting a high relative abundance on the target cells compared to the control strains are colored. Unspecific sybodies are collectively represented by colorless slices. (**c**) Exemplary flow cytometry data of Lp-SG1, Lp-SG4 and Lp-SG6 using propidium iodide for cellular staining (red-channel) and Atto-488-labelled sybody SB400 (green channel). (**d**) Alignment of the major non-conserved region of MOMP and illustration of its location on a homology model of the MOMP monomer. MOMP sequences identical to *Lp-SG6* are framed. (**e**) Cell surface binding of the five identified sybodies to an array of *Legionella* serogroups as quantified by flow cytometry. (**f**) Coomassie-stained SDS-PAGE analysis of MOMP from *Lp-SG6* and Western blot detection via SB400.

To further validate this result, the identified sybodies were individually synthesized, purified, labelled with atto488 and subjected to a flow cytometry screen, using 15 different *Legionella pneumophila* serogroups (Fig. 6d and e) and 52 additional bacterial strains as controls (Supplementary Fig. 9). Interestingly, strong cell-surface binding was observed for *Legionella pneumophila* serogroups 1, 2, 6 and 12, being the only strains with an identical MOMP extracellular region as present in *Lp-SG6* (Fig. 6e), which confirms target-binding in the cellular context. Since binding to purified MOMP is abolished after heat denaturation of the target, the recognized epitope is expected to be three-dimensional (Fig. 6f). In summary, NestLink proved to be highly sensitive and specific for the identification of strongly underrepresented binders, even for extremely challenging targets such as integral membrane proteins embedded in a dense layer of LPS on a Gram-negative pathogen.

## DISCUSSION

NestLink combines advantages of selections and single-clone analyses by processing thousands of pool members as an ensemble, while generating accurate readouts for individual pool members. This allows for direct binder characterization without laborious single-clone handling. We show the potential of the technology by ranking a sybody pool according to their off-rates, by selecting camelid nanobodies against a membrane transporter in solution, and by mining for rare binders that recognized an outer membrane protein target in the native context of a living Gram-negative pathogen.

NestLink builds on a novel flycode library and the unprecedented process of library nesting in combination with NGS, LC-MS/MS and gene synthesis. Unlike previously established NGS and LC-MS/MS based technologies, NestLink benefits from a large, controllable number of unambiguously assignable peptides per binder with optimal detection efficiencies. Remarkably, throughput and cost limitations for NestLink are balanced and none of the technologies was identified as a single major bottleneck. Illumina MiSeq has a throughput of approximately 20 Mio reads, suggesting that 400,000 flycodes can be sequenced per run at an average read redundancy of 50-fold. This corresponds to maximally 13,000-20,000 binders per nested library, each coupled to 20-30 flycodes on average. Depending on the enrichment levels of favorable binders in the input pool and the stringency of the applied selection pressures, only a small subset of available flycodes is subjected to mass spectrometry. Current LC-MS/MS setups can detect several tens of thousands of peptides, corresponding to a few thousand well detected binders per run. Importantly, LC-MS/MS gains in sensitivity upon reduction of sample complexity. Hence, if an input pool is particularly challenging and contains only a tiny fraction of useful binders able to pass the relevant NestLink selection pressures, LC-MS/MS detects them particularly well. Thus, NestLink is not only suitable to uncover large numbers of binder families (see application II), it is also ideally suited for the identification of rare binders, as demonstrated by application III. Of note, binder numbers in NestLink refer to unique binder sequences and are thus not merely analogous to the throughput of single-clone analyses, where rare binders can only be identified upon redundant analysis of many identical clones of enriched binders.

Using NestLink, entire binder pools are processed analogous to the handling of single clones. Hence, it is conceivable to parallelize NestLink and perform selections in screen-like setups in a scalable fashion. Notably, there are massive economic incentives to render deep sequencing, LC-MS/MS and gene synthesis even more efficient in the near future, from which the NestLink method will greatly benefit.

We wish to stress, that the presented technological principle is not restricted to binders. Rather, it may be applicable to any protein pool analysis that permits a spatial separation of desired from undesired library members. NestLink-type approaches may for example allow efficient identification of flycode-tagged, thermostable G protein-coupled receptor mutants with favorable SEC elution profiles, which are suitable for *in vitro* drug screening and structural biology. Similarly, antibodies or enzymes with improved stability and aggregation propensities may be identified.

Since NestLink selections are performed in the absence of a physical genotype-phenotype linkage, large display particles are no longer preventing size-dependent characterization of binder pools in tissues or *in vivo*. Hence, NestLink opens avenues to monitor biodistribution, tissue penetration, immunogenicity or serum half-life for thousands of biopharmaceutical drug candidates at once in a single disease-relevant model organism.

## ACKNOWLEDGMENTS

We thank Dr. Olga Schubert, Dr. Roger Dawson and Dr. Eric Geertsma for their helpful comments on the manuscript and Saša Štefanić for alpaca immunizations. This work was funded by a grant of the Commission for Technology and Innovation CTI (16003.1 PFLS-LS, to MAS), a SNSF Professorship of the Swiss National Science Foundation (PP00P3_144823, to MAS.) and a SNSF BRIDGE proof of concept grant (20B1-1_175192, to PE).

## AUTHOR CONTRIBUTIONS

PE, IZ and MAS developed the conceptual basis of NestLink. PE, BR, CP and MAS designed the flycode library and PE and IZ generated it. PE performed library nesting and selections. NGS was carried out by LO, LP and PE. IZ, FMA, CAJH, DM, HAK and PE performed single-clone analyses via SPR or flow-cytometry. LC-MS/MS was performed and the data was analyzed by PE, BR and CP. PE and MAS wrote the manuscript.

## ONLINE METHODS

### Flycode library design

A random experiment was conducted to simulate and visualize *in silico* the LC-MS/MS detection characteristics of a large number of flycodes using the R environment and the protViz package (https://CRAN.R-project.org/package=protViz^14^. The scripts yield a graphical and numerical output enabling characterization of flycode library dispersity in reversed-phase chromatography and mass spectroscopy. Testing a large number of different sets of input parameters defining flycode lengths, composition of randomized regions and flanking patterns, an optimal flycode library of the sequence “GSX7WZ0-4R” was identified. “GS” corresponds to the C-terminal remainder of a thrombin cleavage site, which enables proteolytic separation from the library of interest. “X7“ corresponds to 7 amino acid positions that encode the following amino acids at their respective frequencies: A: 18 %; S: 6 %, T: 12 %, N: 1 %, Q: 1 %, D: 11 %, E: 11 %, V: 12 %, L: 2 %, F: 1 %, Y: 4 %, W: 1 %, G: 8 %, P: 12%. The constant amino acid “W” was chosen to increase the overall hydrophobicity to the optimal reverse-phase separation range and since a constant amino acid was required at this position for cloning purposes (BfuAI-site). “Z_0-4_” corresponds to the 5 different combinations “no amino acid”, “L”, “QS”, “LTV” or “QEGG”. The C-terminal “R” was chosen because i) its guanidine group allows for an optimal positive charge stabilization and ii) as it enables efficient separation of the flycode from its C-terminal His-tag by trypsin.

Overall, the flycode library was designed to achieve an even spread of hydrophobicity covering the entire range of typical reverse-phase chromatographic separation powers and an optimal m/z - dispersity that falls within the ideal detection window of high-field orbitraps. All randomized regions are devoid of positively charged residues (K, R, H), such that the N-terminus and the C-terminal arginine render each flycode a well-defined doubly-charged species, which is detectable in the ideal m/z range. We confirmed this assumption also for the gas-phase experimentally and found that more than 99 % of flycode precursor ions correspond to doubly charge species. The omission of positively charged residues is also critical in order to render trypsin a site-specific protease (removal of His-tag, see previous paragraph). Methionine and cysteines were omitted to minimize oxidation events, such as cross-linking via disulfide bonds. Glutamate and aspartate are frequent within the randomized stretch to achieve high library solubility at neutral pH, while still allowing efficient reverse-phase binding in the absence of the negative charge at pH 2 (LC-MS/MS conditions). The flycode library exhibits a theoretical diversity of 5.3 x 10^8^.

### Flycode library generation

The flycode library (Fig. 3a) was generated on the basis of the periplasmic expression vector pSb_init (Zimmermann et al., under review) by standard molecular biology techniques and is designated pNLx (Supplementary Fig. 10). Five vector variants were constructed, designated pNLx-pre1, pNLx-pre2, pNLx-pre3, pNLx-pre4, pNLx-pre5, each encoding one of the five flycode C-terminal sequences, all non-variable regions of pNLx1-5 and two BfuAI-sites for barcode insertion in between the flycode C-terminus and the N-terminal part of the thrombin-site. The oligonucleotide “C GTC ACA TTA ACC TGC TAC TCA AGA GGT AGT nnn nnn nnn nnn nnn nnn nnn TGG CAA GTG CAG GTA TAG AAA CGT” was synthesized using trinucleotides (ELLA Biotech) and encodes the 7 randomized positions at their respective frequencies (see previous section). The flanking sequences allowed for PCR amplification and restriction by BfuAI and thus site-directed insertion into pNLx-pre1, pNLx-pre2, pNLx-pre3, pNLx-pre4, pNLx-pre5 resulting in approximately 2×10^7^ clones per construct. Equal mixing of the five sub-libraries resulted in a library size of approximately 1×10^8^ for pNLx. The library was prepared for library nesting by the excision of *ccdB*, followed by agarose gel purification of the linearized vector backbone and gel extraction (Macherey-Nagel).

### Application I: ranking of binders according to their off-rates

#### Library nesting

A pool of sybodies with a convex CDR was used for this experiment, which was previously enriched for MBP-binders by three rounds of ribosome display against MBP, as previously described (Zimmermann et al., under review). After the third round of ribosome display, the recovered sybody pool was amplified by polymerase chain reaction (PCR) using primers 29 and 30 (supplementary table 1) and Q5 Polymerase (NEB) according to the manufacturer’s standard reaction conditions, followed by purification of the PCR product by agarose gel-electrophoresis and gel extraction (Macherey-Nagel). The PCR-amplified sybody pool was sub-cloned via BspQI restriction into the FX cloning vector pINITIAL^15^ containing a kanamycin resistance cassette. After transformation of *E. coli MC1061* cells and plating on agar plates containing 50 µg/ml kanamycin and 1 % (w/v) glucose, the diversity of the pool was restricted to 1,200 – 1,500 cfu by scraping off and cultivation of the appropriate number of colonies in LB containing 50 µg/ml kanamycin and 1 % (w/v) glucose, followed by DNA isolation (Macherey-Nagel). Subsequently, the diversity restricted pool was excised from pINITIAL by BspQI, followed by agarose gel purification and gel extraction using a kit (Macherey-Nagel). The purified, diversity restricted pool and the linearized, purified flycode library (see above) were nested by ligation using 1 µg of pNLx and 700 ng of sybody pool and 10 Weiss Units of T4 ligase (ThermoFischer) in a reaction volume of 40 µl for 1 h at 37°C followed by heat inactivation for 10 min an 65°C. The ligation mix was subsequently transformed in 2 x 150 µl electro-competent *E. coli MC1061* cells followed by recovery at 37°C in 25 ml SOC medium for 30 min. A small fraction of the recovered cells were distributed on LB-agar plates containing 25 µg/ml chloramphenicol for diversity estimation and larger fractions of varying sizes were used to inoculate several 250 ml over-night LB cultures containing 25 µg/ml chloramphenicol. Based on the cfu dilution series on plates, an over-night culture was inoculated with approximately 12,000 – 15,000 cfu and used for DNA Midi preparation (Macherey-Nagel) and the production of a glycerol stock for storage at −80°C.

#### Illumina MiSeq sequencing and flycode assignment

The Illumina MiSeq NGS template was prepared as follows: 100 units of SfiI (NEB) were used to digest 25 µg of the prepared pNLx (containing the nested library) in a reaction volume of 200 µl at 50°C for 1.5 h followed by SfiI inactivation by addition of 8 µl of 0.5 M EDTA. The excised linear nested library was isolated by agarose gel purification followed by gel extraction (Macherey-Nagel). Subsequently, 2×332 ng of double-stranded Illumina adaptor oligonucleotides containing compatible sticky ends (previously generated by DNA-synthesis, supplementary table 1) were site-specifically ligated to both ends of the linearized nested library (600 ng), thereby avoiding PCR amplification (Supplementary Fig. 3). The ligation product containing two adaptors attached was isolated by agarose gel purification, followed by gel extraction (Macherey-Nagel). The concentration of the ligation product was determined by a NanoDrop 2000c Photospectrometer (Thermo Scientific). The ligation product was mixed with differentially indexed ligation products (originating from unrelated experiments for multiplexed Illumina analysis), to obtain an approximate read redundancy of 50 per flycode for each index, aiming for a total of 10 Mio reads. The molarity of the NGS template mixture was subsequently confirmed using the Tapestation 2200 (Agilent) and adjusted to 4 nM. HT1 hybridization buffer was subsequently used to further dilute the library pool. To generate the sample for clustering, 420 µl of the library at 8 pM was mixed to 180 µl of PHiX (Illumina) at 12.5 pM. The sample was sequenced using a 600cycle v3 Miseq reagent kit for 2 x 300 bp paired-end reads on an Illumina MiSeq Sequencer. 8.4 Gb of data were obtained from a single run with >98% reads passing filters, i.e. 14 Mio passed filter reads that had a mean quality score of 35.

For the relevant index of this experiment, 729,932 raw read pairs were obtained and subsequently preprocessed using Trimmomatic (v0.33, parameters: AVGQUAL:20 MINLEN:100) and Flexbar (v2.5, parameters: --pre-trim-left 4 --pre-trim-right 4). 690,066 high quality read pairs were combined using Flash (v1.2.11, default parameters). 618,049 combined reads were obtained, followed by filtering for read length (611,320 reads were ±5% around observed median read length), for flanking patterns of the flycodes (603,488), for flanking patterns of the sybodies (603,084), for reads without N’s (586,643), for the expected construct lengths (492,580), for sequences without stop codons (484,586) and for sequences with correct flycode endings (482,305). The number of unique flycodes was subsequently determined to be 13,620 (minimum of 4 reads per flycode). For each of the 13,620 unique flycodes, a consensus of all binders linked to the same flycode sequence (35 binder sequences on average, due to 35-fold read redundancy) was formed at the amino acid (aa) level and scored for filtering. For each aa position of a binder, the relative fraction of the most frequent aa was calculated as follows: #most frequent aa/(#most frequent aa + #second most frequent aa). The consensus score of a binder corresponds to the average relative fraction over all its amino acid positions. Removing flycodes with a consensus score below 0.9 resulted in 12,160 unique flycodes passing the filter, which were linked to 1,070 distinct binders. On average, 451 sequences (passing all filtering criteria) were obtained per unique binder. An end-pairing overlap of 62 – 68 bp (depending on the flycode length) allowed acquisition of full-length sybody sequences and not merely CDRs. Based on the NGS analysis, a database for MS/MS ion searches (p1875_db8 (release 2016-07-11) was constructed, which assigns each unique binder sequence (identifier) to a virtual protein consisting of its concatenated unique flycodes (Supplementary Fig. 2).

#### Expression of nested library and selections

In order to express the nested library in the vector pNLx in *E. coli MC1061*, the previously generated glycerol stock (see above) was used to inoculate a 37°C overnight pre-culture containing LB and 1 % (w/v) glucose. 2 x 12 ml of saturated pre-culture was used to inoculate 2 x 600 ml of TB containing 25 µg/ml chloramphenicol, followed by induction at an OD_600_ of 0.6 using 0.05 % (w/v) arabinose for 14 h at 20°C. Cells were harvested by spinning at 5,000 g for 15 min, followed by resuspension in 60 ml of TBS (20 mM Tris-HCl pH 7.5, 150 mM NaCl), 10 mM imidazole pH 8 and a pinch of DNase1. The resuspended cells were disrupted using a microfluidizer processor (Microfluidics) at 30,000 lb/in^2^ and the debris was pelleted by centrifugation at 4,400 g for 30 min. The supernatant was loaded on a 1.5 ml Ni-NTA column (Qiagen), the column was washed by 30 ml of TBS containing 30 mM imidazole pH 8, followed by 3 x 2 ml elution using TBS containing 300 mM imidazole pH 8. The eluted nested library was filtered (0.2 µm syringe filter) and was subjected to a SEC run on an Aekta Purifier (GE-Healthcare) system using a HiLoad 16/600 Superdex 200 pg (GE-Healthcare). The nested library members, corresponding to the monomeric binders, were collected and concentrated using centrifugal filters with a 3 kDa cut-off (Amicon Ultra-15) to an absorbance (280 nm) of 2.0. The biotinylated target MBP-biotin at a concentration of 204 µM was prepared as previously described (Zimmermann et al., under review). Three analogous SEC runs were performed in TBS on a Superdex 200 10/300 (GE-Healthcare). The first sample contained 175 µl of the nested library and 60 µl TBS, the second sample contained 175 µl of the nested library and 60 µl of MBP-biotin and the third sample contained 175 µl TBS and 60 µl of MBP-biotin. Superposition of the three runs allowed collecting those early eluting fractions (3 ml total) of the second run, which contained the library members interacting with MBP in solution. The fractions were split into two equivalent 1.5 ml fractions and 150 µl of streptdavidin-sepharose slurry (Thermo Scientific) was added to each fraction, followed by incubation at 4°C for 2 h. The resins were pelleted by centrifugation (swinging-bucket) for 10 min at 200 g and transferred to two Mini Bio-Spin® Chromatography Columns (Bio-Rad: #732-6207). The columns were drained by centrifugation at 50 g for 5 sec in a table-top centrifuge. Column 1: The resin (75 ul) was resuspended by the addition of 500 µl of TBS containing 10 µM non-biotinylated MBP and incubated for 195 seconds (off-rate selection), followed by draining (5 sec at 50 g). Column 2: was not challenged by MBP but otherwise treated identical to column 1. Both columns were washed immediately by 500 µl TBS.

#### Flycode isolation and LC-MS/MS

5 µl of a control binder attached to 28 different flycodes of known sequence (NB-control, see below) at an absorbance of 0.05 (280 nm) was added to both columns as an LC-MS/MS standard. The resins were resuspended in 100 µl of buffer TH (20 mM Triethylammonium bicarbonate (TEAB) pH 8.5, 150 mM NaCl, 2.5 mM CaCl_2_) containing 2.4 units of thrombin (Novagen, #69671-3) and incubated over night at 20°C. The His-tagged flycodes were eluted by centrifugation at 100 g for 10 sec, followed by washing with 2 x 300 µl buffer TH. The flycodes were subsequently pulled down by incubation for 1 h at 20°C with 80 µl Ni-NTA slurry (Qiagen), followed by centrifugation at 800 g for 5 min. The resins were transferred to Mini Bio-Spin^®^ Chromatography Columns (Bio-Rad), washed with 500 µl of buffer TRY (50 mM TEAB pH 8.5, 50 mM NaCl, 2.5 mM CaCl_2_) and subsequently resuspended in 100 µl buffer TRY containing 0.5 µg trypsin (Promega, #V5113), followed by incubation over night at 37°C. The flycode mixture (severed from His-tags) was eluted from the columns by centrifugation for 30 sec at 100 g, followed by a 100 µl wash (50 mM TEAB pH 8.5, 150 mM NaCl) and addition of 20 µl of 5% (v/v) TFA and 200 µl of 3 % (v/v) ACN, 0.1 % (v/v) TFA to the elution. The eluted flycode mixture was loaded onto ZipTips (Millipore, #ZTC185960) pre-treated by washing with 200 µl of 60 % (v/v) acetonitrile (ACN), 0.1 % (v/v) trifluoroacetic acid (TFA), 200 µl of methanol and 200 µl of 3 % (v/v) ACN, 0.1 % (v/v) TFA. The ZipTips were washed by 200 µl of 3 % (v/v) ACN, 0.1 % (v/v) TFA, followed by elution with 2 x 40 µl of 60 % (v/v) ACN, 0.1 % (v/v) TFA and lyophilization of the elution and re-solubilization of the flycodes in 15 µl of 3 % (v/v) ACN, 0.1 % (v/v) formic acid (FA). For application I, flycodes were analyzed using an Easy-nLC 1000 HPLC system operating in single column mode coupled to an Orbitrap Fusion mass spectrometer (Thermo Scientific). 2 µl of the resuspended flycode solution was injected onto an in-house made capillary column packed with reverse-phase material (ReproSil-Pur 120 C18-AQ, 1.9 µm; column dimension 150 mm x 0.075 mm, Temp. 50°C). The column was equilibrated with solvent A (0.1 % formic acid (FA) in water) and peptides were eluted with a flow rate of 0.3 µl/min using the following gradient: 5 - 20 % solvent B (0.1 % FA in ACN) in 60 min, 20 - 97 % solvent B in 10 min. High accuracy mass spectra were acquired with an Orbitrap Fusion mass spectrometer (Thermo Scientific) using the following parameter: scan range of 300-1500 m/z, AGC-target of 5e5, resolution of 120,000 (at m/z 200), and a maximum injection time of 100 ms. Data-dependent MS/MS spectra were recorded in rapid scan mode in the linear ion trap using quadrupole isolation (1.6 m/z window), AGC target of 1e4, 35 ms maximum injection time, HCD-fragmentation with 30 % collision energy, a maximum cycle time of 3 sec, and all available parallelizable time was enabled. Mono isotopic precursor signals were selected for MS/MS with charge states between 2 and 6 and a minimum signal intensity of 5e4. Dynamic exclusion was set to 25 sec and an exclusion window of 10 ppm. After data collection peak lists were generated using automated rule based converter control^16^ and Proteome Discoverer 1.4 (Thermo Scientific).

#### LC-MS/MS data analysis

The two LC-MS/MS runs were aligned in Progenesis QI (Nonlinear Dynamics) with an alignment score of 93.1 %, followed by peak picking with an allowed ion charge of +2 to +5. The fragment spectra with a feature rank-threshold of <5 were exported using deisotoping, charge deconvolution and an ion fragment count limit of 1,000. Mascot 2.5 (Matrix Science) was used for flycode identification by a search against database p1875_db8 (release 2016-07-11, generated as described above) concatenated with an in-house built contaminant database (262 common contaminates). Precursor ion mass tolerance was set to 10 ppm and the fragment ion mass tolerance was set to 0.5 Da. In addition, Scaffold (version Scaffold_4.8.4, Proteome Software Inc.) was used to validate MS/MS based peptide and protein identifications. Peptide identifications were filtered for FDR less than 0.1% by the Peptide Prophet algorithm^17^ and protein identifications were filtered for FDR less than 1.0% containing at least 2 identified peptides. Protein probabilities were assigned by the Protein Prophet algorithm^18^. Proteins that contained similar peptides and could not be differentiated based on MS/MS analysis alone were grouped to satisfy the principles of parsimony. Proteins sharing significant peptide evidence were grouped into clusters. Scaffold spectrum report was imported into Progenesis QI. The two LC-MS/MS runs were normalized against the spiked reference NB-control (see below) by choosing NB-control as a standard protein (normalization factor = 0.81). The MS1 intensity integrals of all non-conflicting flycode features were summed for each binder. We refer to this sum as “binder abundance”. The ratios between the binder abundances at the two columns were plotted for each individual sybody (Fig. 4b, y-axis).

#### Single-clone verification by SPR

Based on the NestLink data (Fig. 4b), several sybody genes were chosen that appeared to exhibit different interaction strengths according to the off-rate selection analysis. All chosen genes correspond to sybodies that were detected with at least 2 unique flycodes on the columns (112 passed this filter in total). The sybody genes were synthesized (General Biosystems) and subcloned into pSb_init, followed by expression and purification analogous to the nested library, the only difference being supplementation of the SEC buffer by 0.05 % (v/v) Tween-20. Off-rates were determined in this buffer using a ProteOn™ XPR36 Protein Interaction Array System (Bio-Rad) using biotinylated MBP immobilized on a ProteOn™ NLC Sensor Chip to 1,000 response units (RU). 5 different dilutions of the purified sybodies were applied to the chip for 245 seconds (association phase), followed by dissociation phases of varying lengths. The off-rates were derived from Langmuir fits.

### Application II: nanobody selections without target immobilization

#### Immunization of alpaca, phage library preparation and phage display

An alpaca was immunized four times with subcutaneous injections at two week intervals, each time with 200 µg purified TM287/28811 in TBS pH 7.5 containing 0.03% β-DDM. Three additional subcutaneous injections of 200 µg protein were performed at two week intervals. Immunizations of alpacas were performed according to the Swiss animal protection regulations (animal experiment licence nr. 188/2011). The nanobody repertoire of the immunized animal served as input for phage display19 and in a separate experiment for NestLink (described in the following paragraphs). Phage 13 display against biotinylated TM287/288 and ELISA screening were performed as previously described (Zimmermann et al., under review).

#### Library nesting, NGS, expression and purification of nested library

After phage library construction from the B-lymphocytes of the immunized alpaca (without performing phage display), the single-stranded nanobody library was amplified by PCR using Alp-Nb_FX_FW_81 and Alp-Nb_FX_REV_82 (supplementary table 1) and GoTaq G2 DNA polymerase (Promega) according to the manufacturer’s standard reaction conditions, followed by purification of the PCR product by agarose gel-electrophoresis and gel extraction using a kit (Macherey-Nagel). The PCR-amplified nanobody pool was subcloned via BspQI restriction into pINITIAL. Nesting of the nanobody pool with the flycode library was performed as described for application I, but using 3,000–4,000 cfu of the nanobody pool (pINITIAL) and 60,000 – 80,000 cfu of pNLx after nesting. NGS was performed as described for application I, but with a consensus score cut-off of 0.99. After filtering 59,974 flycodes linked to 3,390 unique nanobodies were obtained, which were used for the generation of the flycode assignment table (p1875_db10 (release 2017-08-18)).

The nested library was expressed and purified as described above, but using 1.5-fold the culture size and two instead of one 1.5 ml Ni-NTA (Qiagen) columns with all buffer volumes adjusted accordingly. Two runs of SEC (HiLoad 16/600 Superdex 200 pg (GE-Healthcare)) were performed to isolate monomeric nested library members, yielding 20 ml solution at an approximate nanobody concentration of 22 µM, assuming an average molar extinction coefficient of the nested library of 30,000 M^-1^cm^-1^.

#### Pool-internal competition binding experiment

Complex formation using the purified nested library and TM287/288 was performed at three different molar ratios of I) 1:2, II) 31:1 and III) 163:1 in 500 µl TBS containing 0.03 % DDM for 1 h at 4°C. The nested library members that bound to TM287/288 in solution were isolated via separate SEC runs (Superdex 200 10/300 increase (GE-Healthcare)) for the three different molar ratios by collecting the appropriate fractions at the elution volume corresponding to the nanobody-TM287/288 complex. Analogously, three additional SEC runs were performed, each analyzing the purified nested library at one of the three quantities that was used for complex formation (described above) but in the absence of the target. For these background runs, the same fractions (as in the runs with the target) at early elution volumes were collected. Nested library members collected in these background runs represent nanobodies that elute at early elution volumes independent of the target.

#### Flycode isolation, LC-MS/MS and data analysis

Flycodes were individually isolated from 7 different samples: 1) from the purified nested library (200 µl of the monomeric nested library members), 2-4) from the SEC-fractions corresponding to target-nanobody complexes (3 ml of each of the three SEC runs) and 5-7) from the three background SEC runs (3 ml of each of the three SEC runs were collected at the same elution volumes as for the runs isolating target-nanobody complexes). Each sample was spiked with 7 µl of a control binder with 28 known flycodes (NB-control, see below) at an absorbance at 280 nm of 0.052. The 200 µl sample of the purified nested library was diluted to 1.2 ml by TBS for further processing. 100 µl slurry Ni-NTA (Quiagen) was added to each of the 7 different samples, followed by incubation for 2 h at 4°C and pelleting of the resin by centrifugation at 500 g for 5 min. The resins were transferred to Mini Bio-Spin® Chromatography Columns (Bio-Rad: #732-6207) and washed 2 x by 700 µl of buffer Iso (30 % (v/v) isopropanol, 20 mM TEAB, 5 mM imidazole), followed by 3 x 700 µl of buffer TH. The resin was resupsended in 100 µl buffer TH containing 2.4 units of thrombin (Novagen, #69671-3) and incubated over night at room temperature. Subsequently, the resin was washed 5 x by 700 µl buffer TRY containing 10 mM imidazole, followed by elution of the His-tagged flycodes by 2 x 50 µl buffer TRY containing 250 mM imidazole. The eluate was spun through a Microcon filter YM-10 (Amicon, #42407) with a 10 kDa cutoff at 14,000 g at RT. The elution and filtration procedure was repeated by another 2 x 50 µl of the same buffer. Subsequently, 1 µg of trypsin (Promega, #V5113) was added to the flow-through, followed by incubation over night at 37°C. 20 µl of 5 % (v/v) TFA were added to stop the enzymatic digest and the sample was further diluted by addition of 200 µl of 3 % (v/v) ACN, 0.1 % (v/v) TFA. The 7 flycode mixtures were processed by ZipTips (Millipore, #ZTC185960) as described above for application I and analyzed by LC-MS/MS. For application II, flycodes were analyzed by an Easy-nLC 1000 HPLC system operating in trap / elute mode (trap column: Acclaim PepMap 100 C18, 3 µm, 100A, 0.075×20 mm; separation column: EASY-Spray C18, 2 µm, 100A, 0.075×500 mm, Temp: 50°C) coupled to an Orbitrap Fusion mass spectrometer (Thermo Scientific). Trap and separation column were equilibrated with 12 µl and 6 µl solvent A (0.1% FA in water), respectively. 2 µl of the resuspended flycode solution was injected onto the trap column at constant pressure (500 bar) and peptides were eluted with a flow rate of 0.3 µl/min using the following gradient: 5 - 20 % B (0.1 % FA in ACN) in 60 min, 20 - 97 % B in 10 min. High accuracy mass spectra were acquired with an Orbitrap Fusion mass spectrometer (Thermo Scientific) using the following parameter: scan range of 300-1500 m/z, AGC-target of 5e5, resolution of 120,000 (at m/z 200), and a maximum injection time of 100 ms. Data-dependent MS/MS spectra were recorded in the linear ion trap using quadrupole isolation (1.6 m/z window), AGC target of 1e4, 35 ms maximum injection time, HCD-fragmentation with 30 % collision energy, a maximum cycle time of 3 sec, and all available parallelizable time was enabled. Mono isotopic precursor signals were selected for MS/MS with charge states between 2 and 6 and a minimum signal intensity of 5e4. Dynamic exclusion was set to 25 sec and an exclusion window of 10 ppm was used. After data collection, peak lists were generated using automated rule based converter control^16^ and Proteome Discoverer 1.4 (Thermo Scientific). Two technical replicates were recorded for each sample (total of 14 LC-MS/MS runs).

Two alignments were generated using Progenesis QI: 1) the two LC-MS/MS replicates of the sample representing the purified, monomeric nested library members (alignment score of 94.5 %) and 2) LC-MS/MS runs of the samples corresponding to the collected fractions of the pool-internal competition experiments or their respective background runs (alignment scores between 69.0 and 97.6 %). Note that aligning LC-MS/MS runs was not per se necessary for the NestLink data analysis performed here, but it allowed parallel workflows for similar recordings and thus facilitated data processing in Progenesis QI. Peak picking, peptide filtering and peptide exporting was performed as described for application I (see above). Mascot 2.5 (Matrix Science) was used for flycode identification by two searches (one search for each alignment) against 3 databases per search i) p1875_db10 (release 2017-08-18, generated as described above), ii) p1875_db8 (release 2016-07-11) both have been concatenated with an in-house built contaminant database, and iii) Swissprot database (release 20140403) concatenated with its decoyed entries. Mascot search parameters and processing in Scaffold were analogous to application I (described above). After re-import into Progenesis QI, the LC-MS/MS runs were normalized using the spiked standard NB-control. A normalization factor of 1.00 was obtained for the first alignment and factors between 0.51 – 1.15 were obtained for the second alignment. Note that, normalization was not essential for the analysis performed here, but it served as a control, since extreme normalization factors would hint at inconsistencies in sample preparation. The MS1 intensity integrals of all non-conflicting flycode features were summed for each nanobody in each sample (binder abundance). Binder abundances were averaged between the two technical LC-MS/MS replicates per sample (see above). The “relative abundance” corresponds to the fraction of an individual nanobody abundance relative to the total of all nanobody abundances detected in the same LC-MS/MS run (100 %), thus calculating the relative abundance corresponds to a sample-internal normalization.

Nanobody sequences exhibiting an increase in relative abundance in the following order, 1) purified nested library (input pool), 2) SECI, 3) SEC I, 4) SECIII (target-bound fractions for 2)-4)) and at least 4 detected flycodes on SEC were considered as strong binder hits. Their sequences were extracted from the NGS database and their CDR3 regions were aligned using the alignment tool of the software CLC (Qiagen), followed by editing in Jalview (Fig. 5c). Only nanobodies exhibiting more than 10-fold higher binder abundances in the complex runs compared to the same “shifted” fraction of the background runs (no target, see above), were included in the analysis.

#### Single-clone verification by SPR

Genes of 11 “binding-nanobodies” (see above alignment) and 4 “negative-control-nanobodies” (detected in the purified nested library, but not at the target) were synthesized (General Biosystems) and subcloned into pSb_init, followed by expression and purification analogous to the nested library. Binding kinetics were determined using a ProteOn™ XPR36 Protein Interaction Array System (Bio-Rad) using biotinylated TM287/288 immobilized on a ProteOn™ NLC Sensor Chip to 1,500 response units (RU) and TBS supplemented with 0.03 % (w/v) DDM. An initial SPR screen was performed at a single concentration of 100 nM for each nanobody. For the “binding-nanobodies”, this screen revealed that two nanobodies (NL2.1 and NL11.1) exhibited off-rates that were too slow to be determined by the ProteOnTM (< 5E-5 s-1) and two nanobodies exhibited significantly higher dissociation constants than 100 nM (NL1.3 and NL7.1). From the 11 purified “binding-nanobodies”, 7 were therefore used for accurate determination of kinetic parameters. To this end, 5 different dilutions of the purified nanobodies were applied to the chip for 245 seconds (association phase), followed by dissociation phases of varying lengths. The data were fitted using the Langmuir method. None of the 4 “negative-control-nanobodies” exhibited a binding signal.

### Application III: specific recognition of an outer membrane protein in the cellular context

#### Purification of the major outer membrane protein (MOMP) from L. pneumophila serogroup 6

*L. pneumophila serogroup 6* strain DSM25182 was grown at 37 °C on BCYE agar (BBL BCYE Agar Base, BD). Single colonies were inoculated in 5 ml liquid BCYE media (Legionella BCYE Growth Supplement, VWR) and grown to stationary phase by shaking overnight at 37 °C. 0.5 ml of the densely grown BYE pre-culture was used to inoculate 500 ml cultures of liquid BYE media (10 g/l yeast extract, 0.25 g/l ferric pyrophosphate, 1 g/l α-ketoglutarate, 0.4 g/l L-cysteine, 7.2 g/l ACES buffer adjusted to pH 6.9) and grown to an OD_600_ of 0.9. Bacteria were harvested by centrifugation at 4,000 g for 10 min, washed in PBS and centrifuged again at 4,000 g for 10 min.

The MOMP protein was purified from a total of 8 liters of BYE culture according to Gabay et al.20. Briefly, the harvested bacteria were resuspended in lysis buffer (0.1 M sodium acetate, pH 4, 0.45 M CaCl2, 0.45 % Zwittergent 3-14, 10 mM beta-mercaptoethanol, 1 mM phenylmethylsulfonyl fluoride), sonicated for 30 s in a sonicating water bath (Elmasonic P) and cooled at 0 °C. Ice-cold absolute ethanol was added dropwise to a final concentration of 20 % ethanol (v/v) and the mixture was stirred for 30 min at room temperature. The preparation was centrifuged at 17,000 g for 10 min and the supernatant was discarded. The pellet was suspended again in lysis buffer and the mixture was sonicated for 30 s using a tip (Branson Sonifier B12), treated with ethanol, and centrifuged at 17,000 g for 10 min. The supernatant, containing MOMP, was collected, treated with ice-cold absolute ethanol to a final concentration of 75 % (v/v) to precipitate proteins, incubated over night at −20 °C and centrifuged at 20,000 g for 35 min. The pellets were suspended in 50 mM Tris-HCl, pH 8.0, 10 mM EDTA, 0.5 % Zwittergent 3-14 and centrifuged at 20,000 g for 35 min to remove insoluble protein. The sample was applied onto two HiTrap FF DEAE columns (GE Healthcare) that were connected in a row and equilibrated with Buffer A (50 mM Tris-HCl, pH 8.0, 10 mM EDTA, 0.05 % Zwittergent 3-14). Bound protein was eluted by applying a 50 ml salt gradient of 0.13 M to 1 M NaCl with Buffer B (50 mM Tris-HCl, pH 8.0, 1 M NaCl, 10 mM EDTA, 0.05 % Zwittergent 3-14) on an Aekta Prime. Elution fractions containing MOMP were pooled and treated with ice-cold absolute ethanol to a final concentration of 75 % (v/v), incubated over night at −20 °C and centrifuged at 20,000 g for 35 min to collect precipitated proteins. The pellet was suspended in a minimal volume of 50 mM Tris-HCl, pH 8.0, 10 mM EDTA, 0.5 % Zwittergent 3-14 and centrifuged at 20,000 g for 35 min before injection onto an S200 10/300 (GE-Healthcare) equilibrated with 10 mM Tris, pH 8, 200 mM NaCl, 10 mM EDTA, 0.05 % Zwittergent 3-14. Eluted fractions were analyzed by SDS-PAGE and fractions containing MOMP were pooled, flash-frozen and stored at −80 °C.

#### Sybody selections against detergent-purified MOMP of Lp-SG6

Purified MOMP of LP-SG6 was biotinylated by EZ-Link™ Sulfo-NHS-LC-LC-Biotin (Thermo Fischer # 21338) at a molar ratio of 1:2.5 at 4°C overnight. Free biotin was removed by SEC in TBS containing 0.03% DDM. The enriched sybody pool was generated using a synthetic nanbody library with a convex CDR3 region (Zimmermann et al., under review). Briefly, sybodies were selected by performing one round of ribosome display followed by two rounds of phage display. qPCR revealed a 1.5 fold enrichment of the convex Sybody pool against MOMP compared to AcrB as negative control. The enriched sybody pool was subcloned into the pSb_init and single clones were picked for small scale expression and ELISA. ELISA revealed approximately 50 % positive hits. 12 % of the hits exhibited specific ELISA signals for detergent solubilized MOMP and showed only background signals against the negative control AcrB.

#### Library nesting, NGS, expression and purification of nested library

Library nesting was performed as described for application I. 1,400 – 1,700 cfu and 20,000 – 26,000 cfu were chosen for diversity restriction of sybodies and nested library members, respectively. A consensus score cut-off of 0.99 was used for NGS data filtering analogous to application II. This resulted in a nested library covering 1,444 unique sybodies and 23,598 unique flycodes (database p1875_db9 (release 2017-01-05)). The nested library was expressed and purified as described for application I. Monomeric nested library members (input for pull-down experiment) were selected by a SEC run on an Aekta Purifier (GE-Healthcare) system using a HiLoad 16/600 Superdex 200 pg (GE-Healthcare).

#### Selection for cell-surface binders by a pull-down experiment

4 x 3 ml of the monomeric nested library members (eluted from SEC) were added to 4 individual test tubes each containing 1 ml of either *Lp-SG6*, *Escherichia coli, Citrobacter freundii* or *Lp-SG3* at an OD_600_ of 50 in TBS at pH 7.5 supplemented with 0.5 % BSA. All subsequent steps, including LC-MS/MS were carried out independently for the 4 samples. After incubation for 5 min, cells were pelleted by 17 centrifugation at 4,000 g for 10 min, followed by resuspension of the cells in 5 ml PBS at pH 7.5. Pelleting and resuspension was repeated twice to remove low affinity sybodies.

#### Flycode isolation, LC-MS/MS and data analysis

The pelleted cells were resuspended in 5ml of 100 mM Tris/HCl (pH 7.5), 750 mM NaCl, 2 % (w/v) n-octyl-β-D-glucopyranoside, 50 mM imidazole pH 8.0 containing a pinch of DNaseI and approximately 1 µg of a control binder with 28 known flycodes (NB-control, see below). After 10 min incubation, 20 ml of 6 M GdmCl were added, followed by incubation for 20 min at 20°C. Insoluble components were pelleted by spinning at 4,400 g for 30 min. The supernatant was filtered (0.2 µm syringe filter), followed by the addition of 100 µl slurry Ni-NTA (Qiagen) to the supernatant of each sample and incubation for 2 h at 4°C. The resin was pelleted by centrifugation at 1,500 g for 30 min. The resin was transferred to Mini Bio-Spin® Chromatography Columns (Bio-Rad) for subsequent flycode isolation analogous to application II (see above). For application III, flycodes were analyzed in dublicate by a Waters M-class UPLC system (Waters AG) operating in trap/elute mode coupled to a Q-Exactive HF mass spectrometer (Thermo Scientific). The LC-system were equilibrated with 99% solvent A (0.1% formic acid (FA) in water) and 1% solvent B (0.1% FA in ACN). Trapping of peptides was performed on a Symmetry C18 trap column (5 µm, 75 µm X 250 mm, Waters AG) at 15 µl/min for 30 sec. Subsequently, the peptides were separated using a HSS T3 C18 reverse-phase column (1.8 µm, 75 µm X 250 mm, Waters AG) and the following gradient: 1-40% B in 60 min; 40-98% B in 5 min. The flow rate was constant 0.3 µl/min and the temperature was controlled at 50°C. High accuracy mass spectra were acquired with a Q-Exactive HF mass spectrometer (Thermo Scientific) that was operated in data dependent acquisition mode. A survey scan was followed by up to 12 MS2 scans. The survey scan was recorded using quadrupole transmission in the mass range of 350-1500 m/z with an AGC target of 3E6, a resolution of 120,000 at 200 m/z and a maximum injection time of 50 ms. All fragment mass spectra were recorded with a resolution of 30,000 at 200 m/z, using quadrupole isolation (1.2 m/z window), an AGC target value of 1E5 and a maximum injection time of 50 ms. The normalized collision energy was set to 28%. Dynamic exclusion was activated and set to 30 sec with a mass tolerance of 10 ppm. After data collection, peak lists were generated using automated rule based converter control^16^ and Proteome Discoverer 1.4 (Thermo Scientific).

Using Progenesis QI, the 8 LC-MS/MS runs of the 4 pull-down samples (2 replicates) were aligned and analyzed as described for application II. Alignment scores of 86.1 - 98.6 % were obtained. Note that aligning LC-MS/MS runs was not per se necessary for the NestLink data analysis performed in application II, but it allowed parallel workflows for similar recordings and thus facilitated data processing in Progenesis QI. Peak picking, peptide filtering and peptide exporting was performed as described for application I (see above). Mascot 2.5 (Matrix Science) was used for flycode identification by a search against database p1875_db9 (release 2017-01-05, generated as described above) concatenated with an in-house built contaminant database. Mascot search parameters and processing in Scaffold were analogous to application I (described above). After re-import into Progenesis QI, the LC-MS/MS runs were normalized using the spiked standard NB-control (normalization factors between 0.97 and 2.34 were obtained). Note that normalization was not essential for the analysis performed here, but it served as a control, since extreme normalization factors would hint at inconsistencies in sample preparation. In analogy to application II, the MS1 intensity integrals of all non-conflicting flycode features were summed for each sybody (binder abundance). The binder abundances were averaged between the two technical LC-MS/MS replicates and each sample was internally normalized by calculating the relative abundance for each sybody.

#### Single-clone verification by flow cytometry

5 sybodies exhibiting flycode coverages of more than 20 %, more than 5 unique flycodes detected and 12 – 100 fold higher relative abundances at Lp-SG6 than at any other strain, were chosen for single-clone analysis by flow cytometry. To this end, the identified sybody genes were synthesized, expressed and purified as described for application I. Subsequently, the sybodies were labelled in the presence of a 1.2 fold molar ratio of AlexaFluor 488-NHS (Alexa488-NHS) and the free dye was removed by dialysis (6,000-8,000 MWCO, Spectra/Por®). Coupling efficiencies were calculated from the absorbance at 280 nm and 488 nm, respectively. The average number of dyes per sybody molecule ranged from 0.8 to 1.1.

For the single-clone verification of cell-surface binding to whole Legionella by flow cytometry, 14 different Legionella pneumophila serogroups and 50 additional bacterial strains were fixed by glutaraldehyde treatment (strains are listed in Supplementary Fig. 9). To this end, Legionella pneumophila strains were grown in buffered yeast extract (BYE) broth (10 g/L yeast extract, supplemented with Legionella BCYE Growth Supplement from VWR) by shaking at 37 °C. Other bacteria were grown in liquid media according to the strain provider’s specifications (DSMZ, NCTC or ATCC). The bacterial cells were washed three times by centrifugation for 10 min at 4,226 g and resuspension in 20 ml PBS per 200 ml bacteria culture. After the last centrifugation, the cells were resuspended in 10 ml PBS containing 2.5 % glutaraldehyde, vortexed and incubated for two hours at room temperature in the dark. The fixed cells were then washed three times in PBS as described above.

Fixed bacterial strains at a concentration of 100,000 cells/ml were incubated with 0.5 µg/ml of Alexa488 labelled single-clone sybody and 0.5 µg/ml propidium iodide for one hour at room temperature. To test for cell-surface binding to bacterial cells, the samples were analyzed by flow cytometry using a CytoFLEX (Beckman Coulter), equipped with a 488 nm laser and filter sets of 525/40 (green channel) and 690/50 (red channel). To reduce background noise, a threshold of 400 was used on the green channel and a threshold of 550 was used on the red channel. The analyses were performed at a flow rate of 100 µl/min.

### Production of control binder for LC-MS/MS run normalization

Using SapI restriction and ligation, a clone of a convex sybody (Sb_MBP#1) encoded in the vector pINITAL was inserted in the flycode library vector via replacement of the negative selection marker ccdB. After transformation of E. coli MC1061 cells and plating on agar plates containing 25 µg/ml chloramphenicol and 1 % (w/v) glucose, 28 cfu were picked for cultivation as a pool in LB containing 25 µg/ml chloramphenicol and 1 % (w/v) glucose, followed by DNA isolation (Macherey-Nagel) and glycerol stock production. The sample was processed by NGS as described above to determine the sequences of the 28 flycodes. The identified flycodes linked to Sb_MBP#1 were concatenated and formatted as an entry of a mascot search database, appropriate for manual addition to any other NestLink database. Sb_MBP#1 linked to its flycodes was expressed in and purified from E. coli MC1061 as described for nested libraries (see application I).

#### Data availability

Mass spectrometry data are available via ProteomeXchange with identifier PXD009301. NGS datasets are available on the European Nucleotide Archive (ENA) under accession number PRJEB25673. The custom software used to design the flycode library and to filter and analyze NGS data is available on request.

---

**Table S1:**
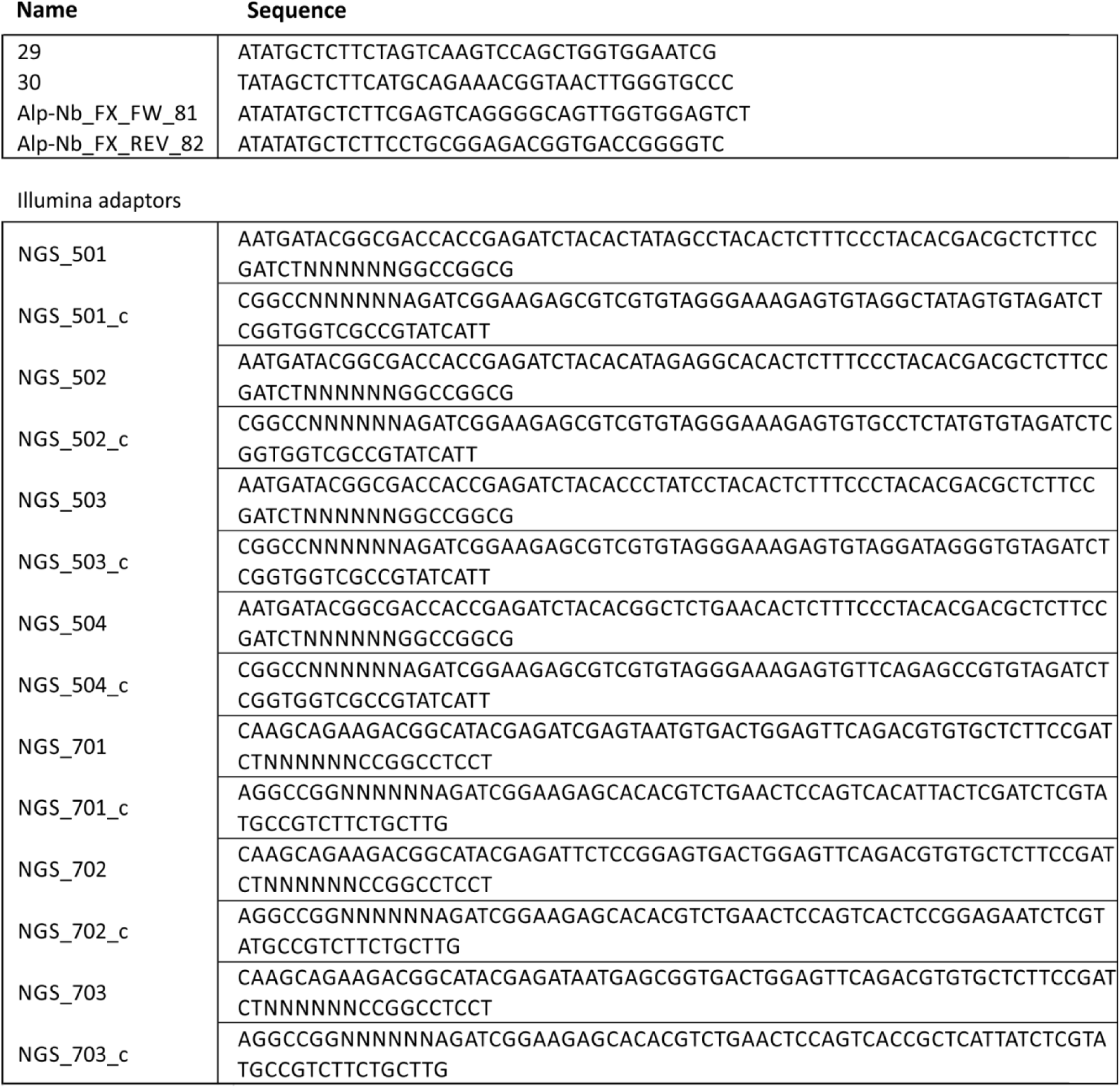
Oligonucleotides used for PCR amplification and Illumina adaptor ligation. Double-stranded Illumina adaptors were generated by annealing of the corresponding complementary oligonucleotides, resulting in sticky ends compatible with the SfiI-excised nested library.

**Figure S1:**
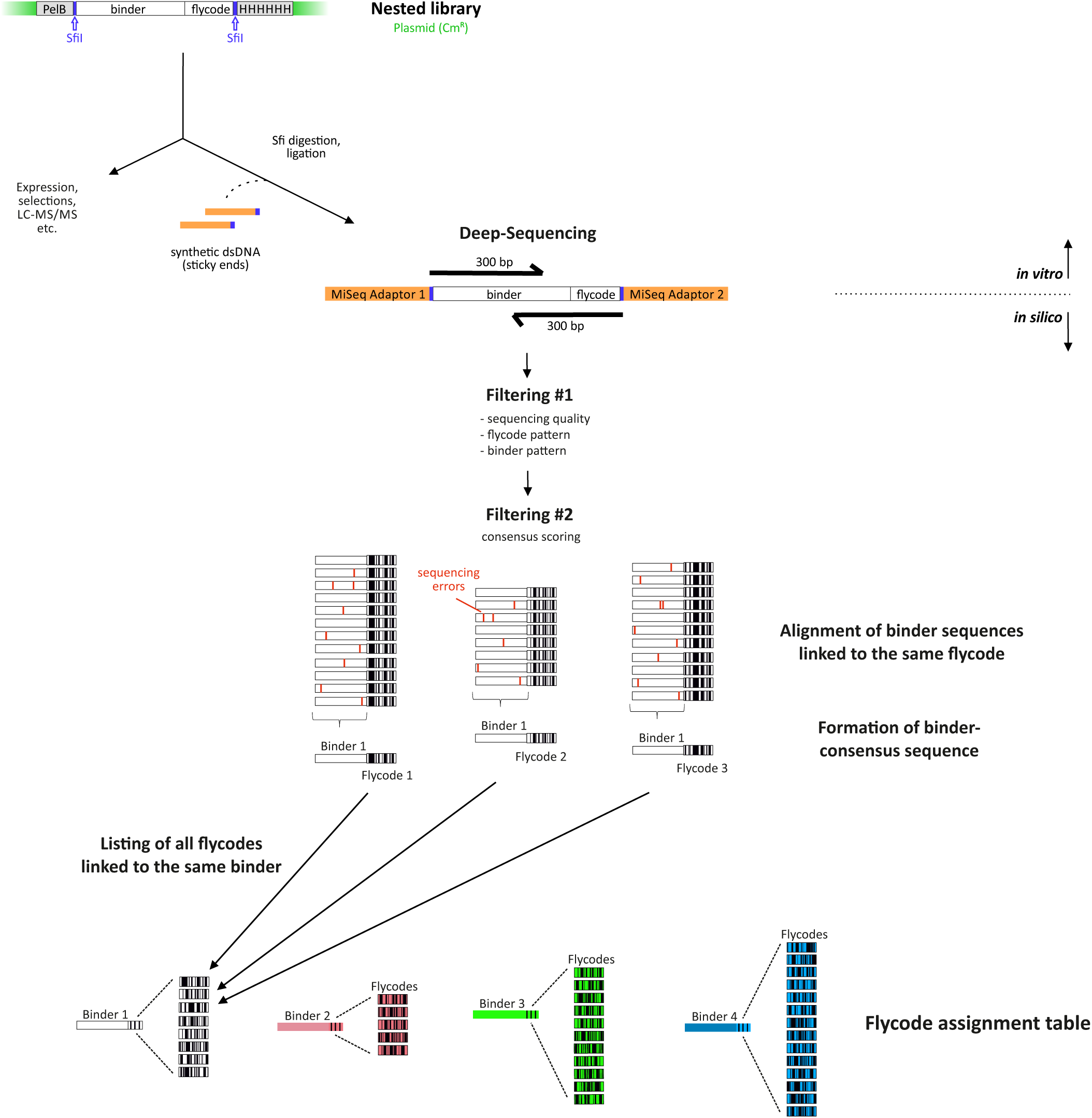
Scheme illustrating NGS and sequence data analysis for NestLink. The nested library is prepared for Illumina MiSeq NGS by a restriction digest with SfiI. This creates two different trinucleotide sticky ends, allowing for site-specific ligation of synthetic, double-stranded MiSeq adaptors containing complementary sticky ends. It is critical to avoid PCR for Illumina adaptor joining, as it introduces a large number of recombination events and inevitably causes the attachment of the same flycodes to several different binders (see Supplementary Fig. 3). By applying Illumina 2×300bp paired-end sequencing to read nested single domain antibody library, a 47-90 bp overlap is achieved (depending on the length of the nested binders and flycodes). Thus, it allows for an accurate acquisition of full-length sybody or nanobody sequences. Indexed adaptors permit for a multiplexed analysis of different nested libraries in a single NGS run. Based on the flycode diversity estimations obtained by cfu counting during library nesting, the concentrations of differentially indexed nested libraries are adjusted to yield about 50 raw-reads per flycode. Raw reads are first filtered for sequencing quality and expected sequence patterns. In a second filtering step, the high read redundancy allows calculations of representative consensus scores for nanobody sequences attached to the same flycode. A stringent consensus score filter is applied, which removes sequencing errors with high efficiency and eliminates rare flycodes attached to more than one binder of the library from further analyses. The NGS output is a database of unique nanobodies and their attached set of flycodes (flycode assignment table).

**Figure S2:**
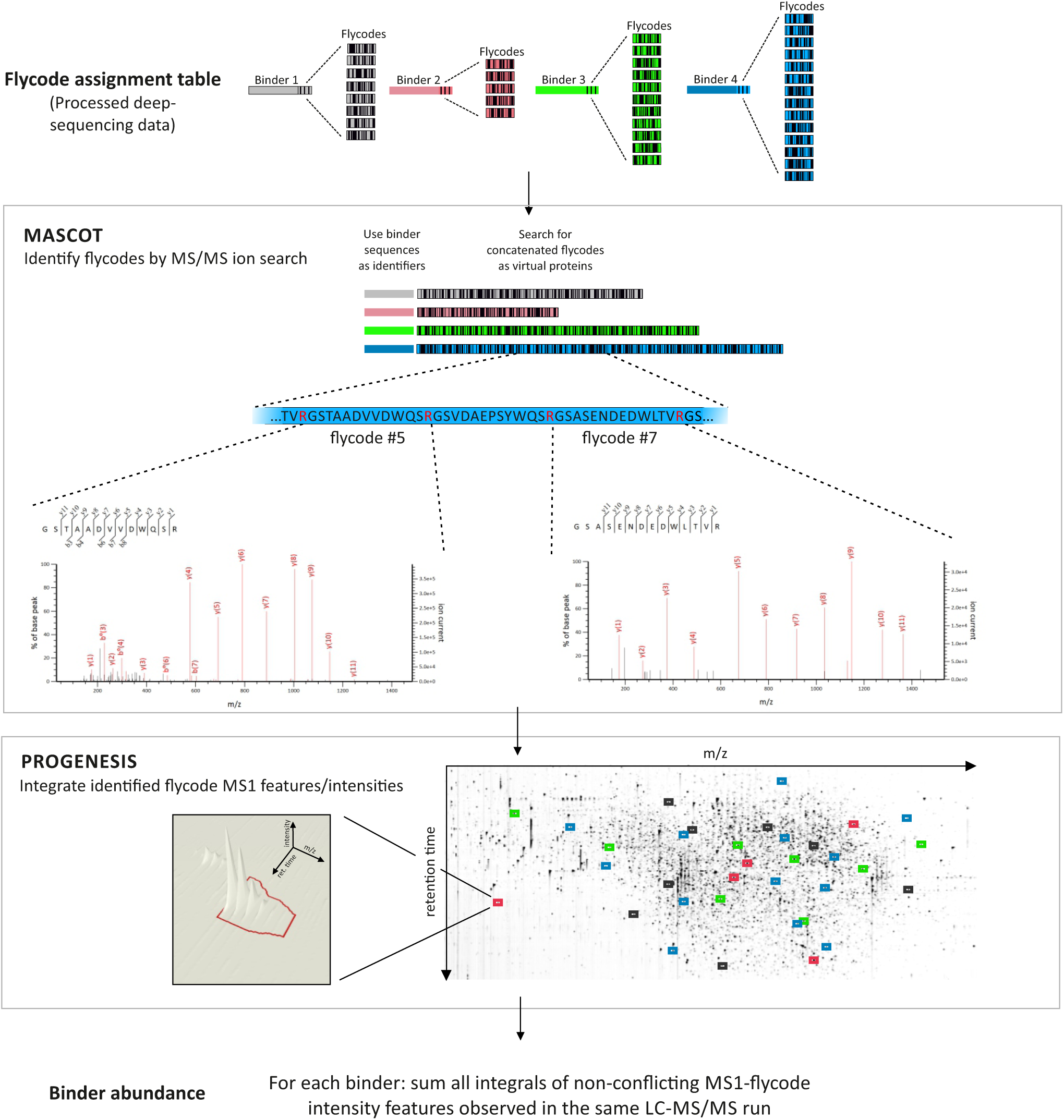
LC-MS/MS analysis for NestLink. The software Mascot15 was used for flycode identification and Progenesis for binder abundance determination. In a first step, the flycode assignment obtained by NGS is converted into a list of concatenated unique flycode sequences (proteins) and identifiers (binder sequences). The concatenated flycodes can be understood as virtual proteins that are searched for by Mascot. To this end Mascot performs first a tryptic digest of the virtual proteins *in silico.* Since flycodes do not encode trypsin cleavage sites internally, the *in silico* digest reverts the concatenation process via cleavage at the flycode terminal arginines. In a second step, the individual flycodes are matched to the recorded MS/MS spectra, followed by the calculation of flycode and binder scores, analogous to peptide and protein scores in proteomics. In a second step, the MS1 precursor ion features of identified flycodes are integrated using the software Progenesis. What we refer to by “binder abundance” corresponds to the sum of all unique (non-conflicting) flycode feature MS1 intensity integrals of a particular binder. After filtering for a minimal number of detected flycodes (see online methods), the experimental outcome for NestLink applications II and III is analyzed by calculating “relative binder abundances”, which refers to the fraction of a particular binder abundance relative to the sum of all binder abundances in a sample (i.e. one LC-MS/MS run). This sample-internal normalization enables to monitor changes of binder frequencies (enrichment or depletion) within pools that have been subjected to different selection pressures.

**Figure S3:**
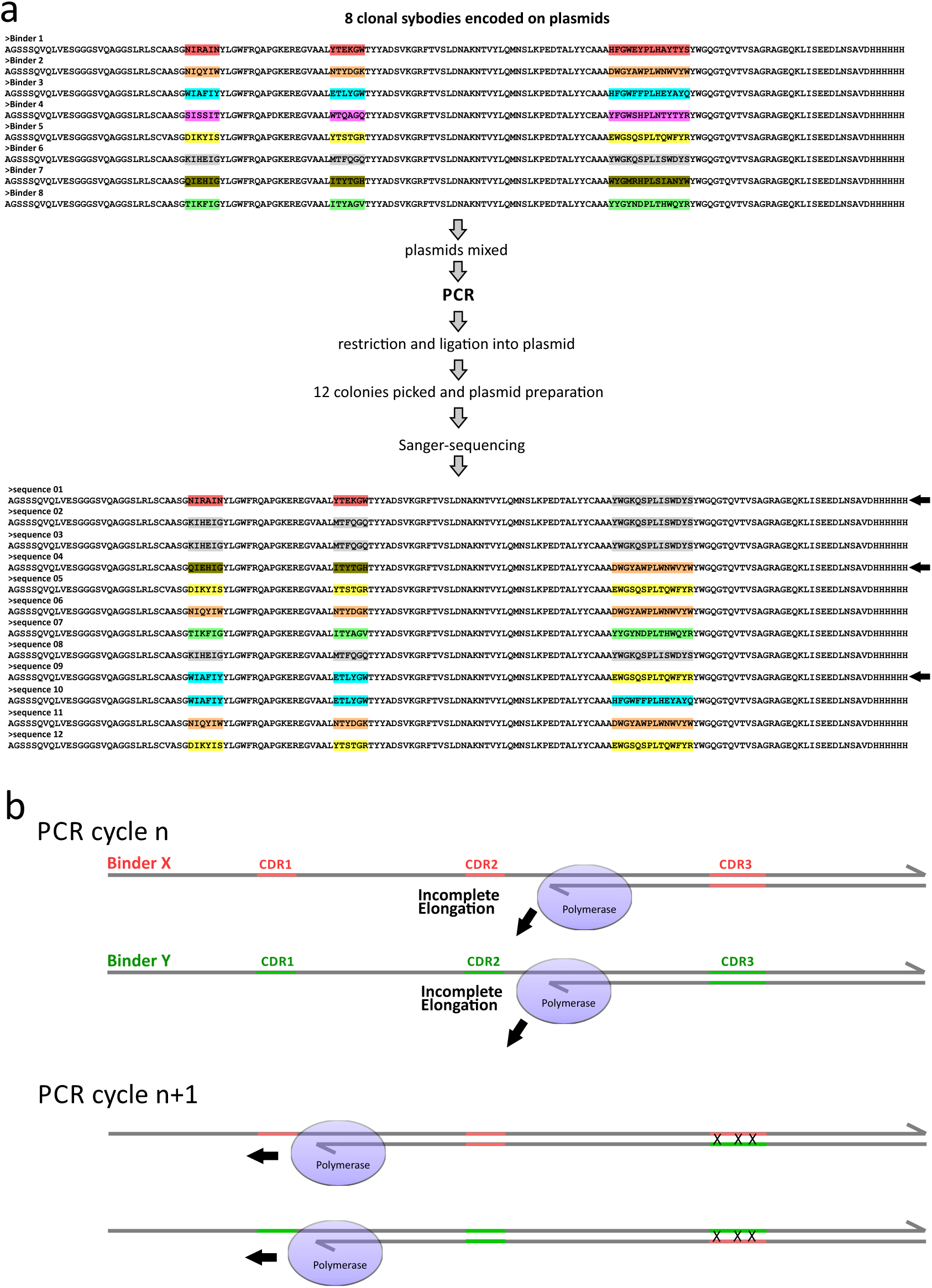
PCR amplifications need to be avoided for library nesting and NGS library preparation. We initially tested a PCR-based protocol for NGS-adaptor attachment. Contrary to what we expected, NGS data analysis revealed that the majority of flycodes were linked redundantly to different binder sequences and could thus not be unambiguously assigned to individual library members. Based on previous reports^16–19^, we assumed that PCR amplification resulted in recombination events. Therefore, we tested this hypothesis experimentally. (**a**) Eight clonal sybodies were pooled and amplified by PCR, followed by sub-cloning into a plasmid and Sanger sequencing of individual clones. Three out of twelve obtained sequences exhibited recombined CDRs. (**b**) Mega-primer formation model. During PCR, incomplete elongation reactions result in mega-primer formation, which can anneal to alternative library members in subsequent cycles and cause recombination events. Since our libraries contain extensive homologous regions, recombination due to mega-primer formation represents a critical problem. Therefore, the NestLink protocol (library nesting and NGS adaptor attachment) operates completely independent of PCR.

**Figure S4:**
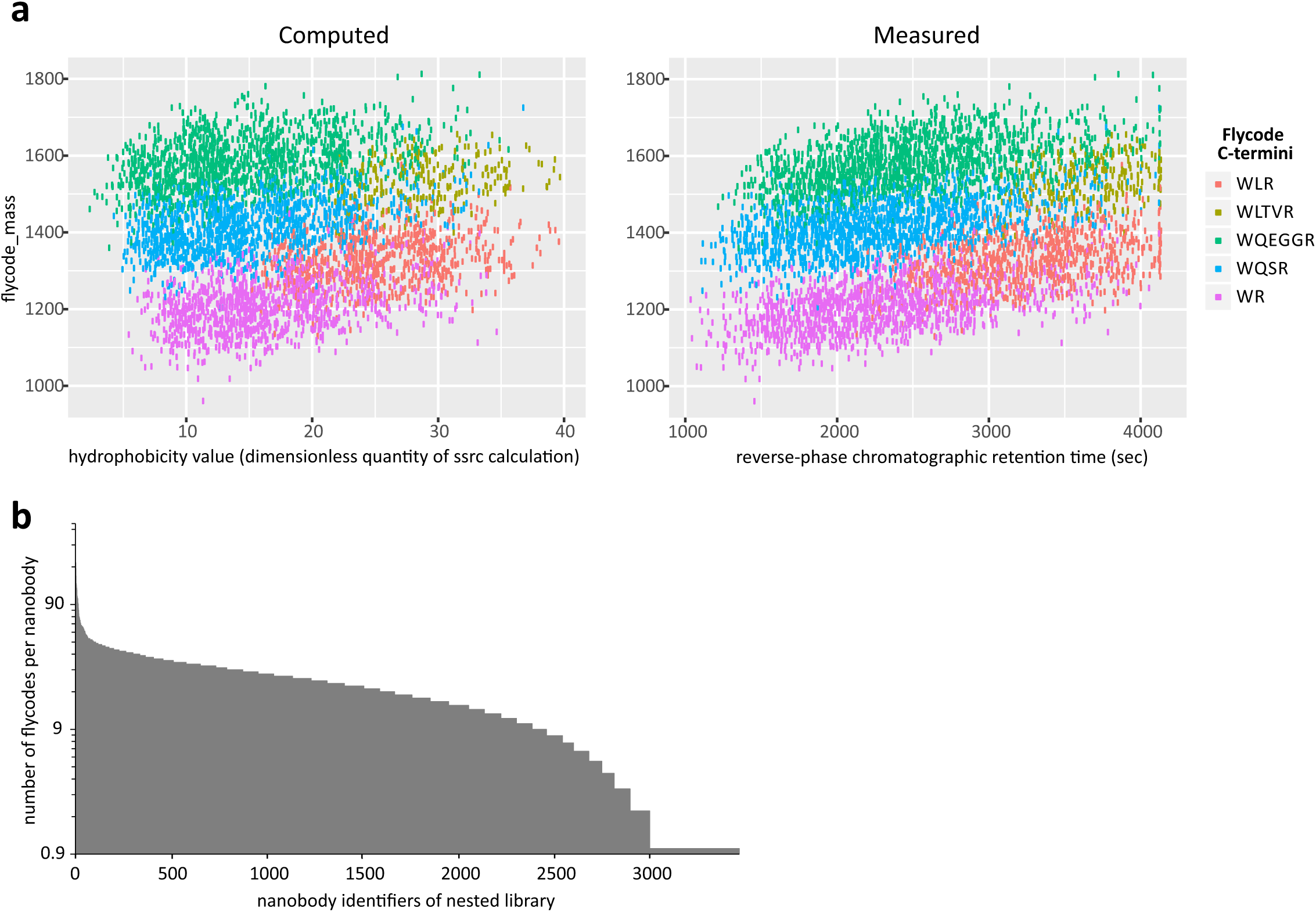
Flycodes harbor five distinct C-terminal sequences to achieve maximal LC-MS/MS dispersion. (**a**) Five distinct flycode C-terminal sequences (red, yellow, green, blue, purple) were designed to cover all areas of the optimal LC-MS/MS detection window with respect to m/z and hydrophobicity. The right panel shows the mass (y-axis) and retention time (x-axis) of 5,202 flycode precursor ions of application II detected by LC-MS/MS (Mascot scores higher than 40). The left panel depicts the simulated dispersion of the same set of flycodes, with hydrophobicities predicted by Sequence Specific Retention Calculator (SSRC)^14^. (**b**) Histogram depicting the number of unique flycodes per nanobody of the nested library used in application II.

**Figure S5:**
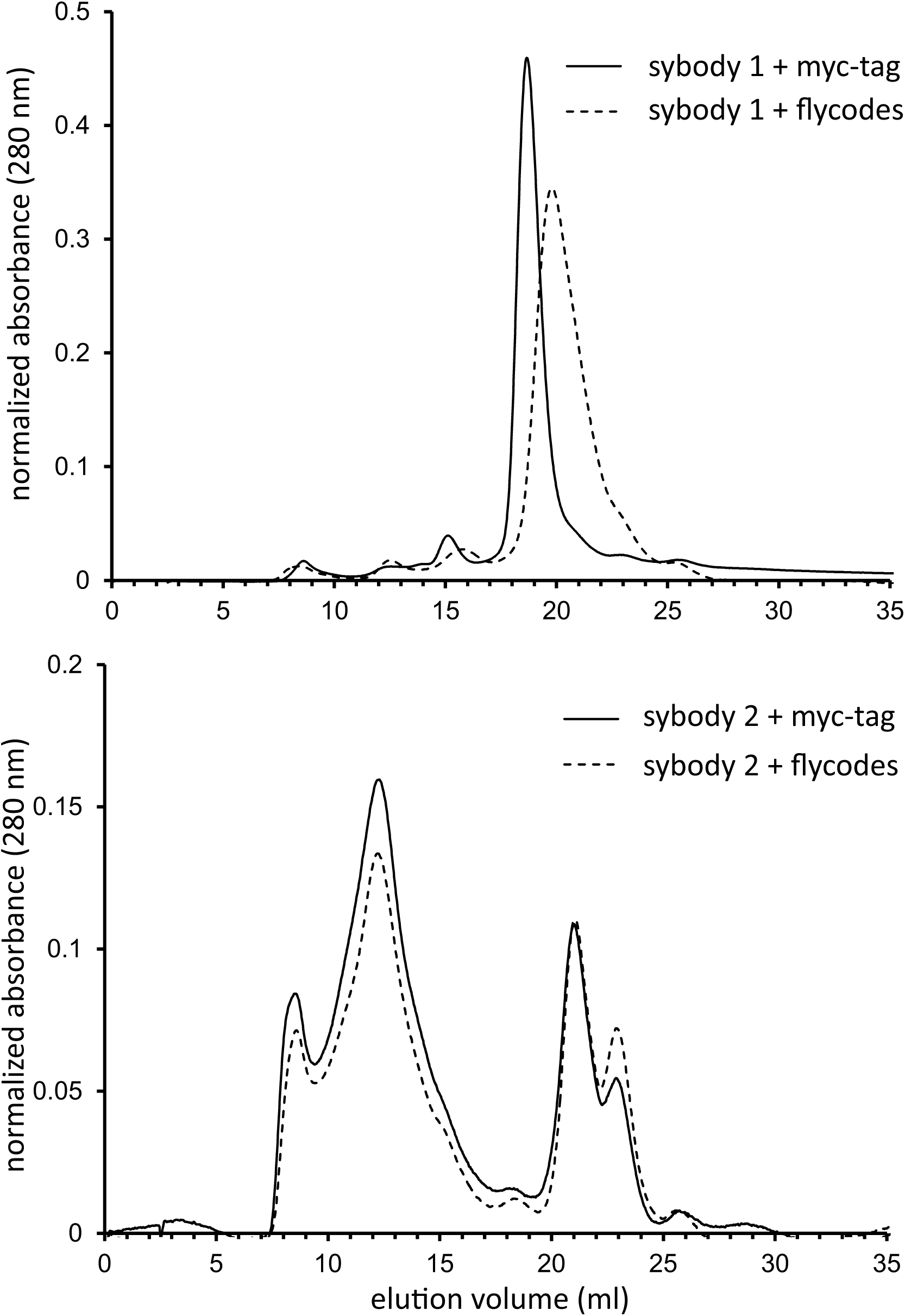
Characterization of a monomeric and an oligomerizing sybody in the presence and absence of flycodes. A monomeric and an oligomerizing sybody were individually fused to more than 1,000 flycodes at the genetic level, followed by expression and purification of the fusion proteins via His-tag. As controls, both binders were expressed and purified without flycodes. Purified proteins were separated by SEC using a Superdex 200 increase 10/300 GL column.

**Figure S6:**
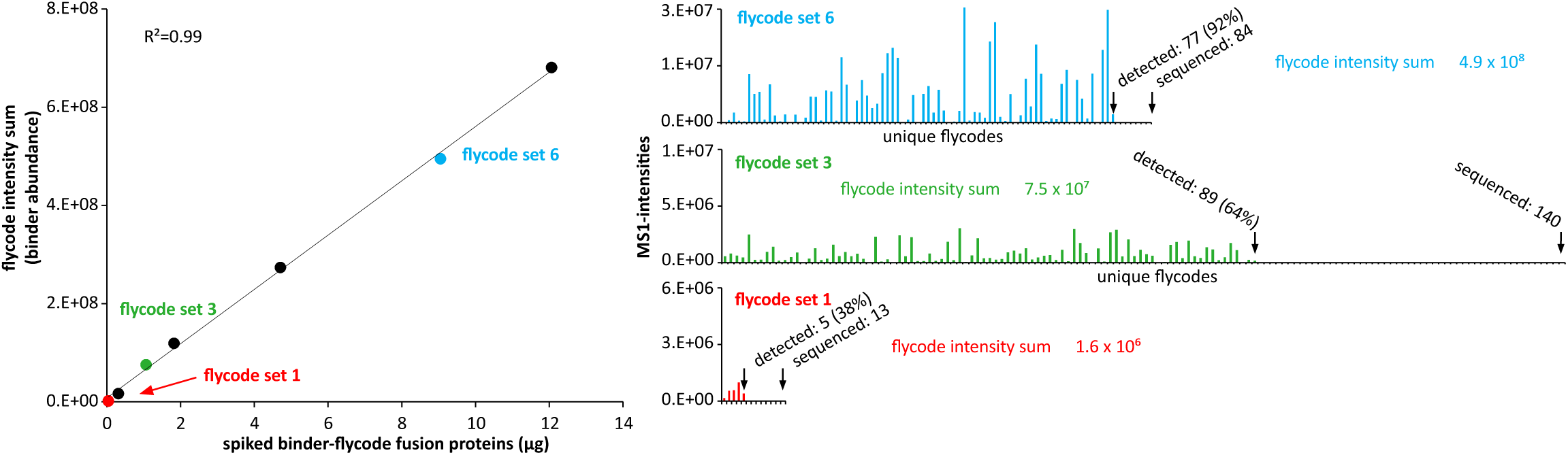
Titration experiment to evaluate flycode detection by LC-MS/MS. One nanobody (identical sequence throughout the experiment) was fused to different sets of known (previously sequenced) flycodes. Different sets of flycodes fused to the nanobody were separately expressed and purified (henceforth and in the graph called “flycode-sets”). After concentration determination of the purified flycode-sets, they were titrated into 50 ml of a bacterial lysate at the concentrations indicated in the main plot (left) on the x-axis. Subsequently, the flycodes were extracted from the lysate as a whole and analyzed by one LC-MS/MS run (2 biological replicates). Subsequently, the MS1 intensity sums over all detected flycodes were calculated for each flycode set, as indicated on the y-axis (left). For three exemplary flycode-sets (red, green, blue) individual flycode MS1 intensities are shown (right). Our interpretation of the experimental outcome is given in the following: First, the current procedure was able to reliably detect less than 50 ng of an individual binder in an inhomogeneous background. Second, more than 90% of the flycodes were detectable for highly concentrated binders, suggesting that NGS and mass spectrometry data are in close agreement. Third, within the investigated range of concentrations spanning 2.8 orders of magnitudes, the flycode intensity sum correlates linearly with the titrated binder quantity. We expect a non-linear response at higher concentrations, due to saturation of ion detectors or limited ionization energy. Fourth, MS1 intensities of individual flycodes within a flycode-set vary strongly, presumably due to differences in their abundance at the gene level, expression, purification, ionization and identification efficiencies. However, if a large number of flycodes belonging to the same flycode-set is detected, the variability is averaged out by summing the corresponding MS1 intensities. Although this experiment implies that flycodes could be used for absolute quantification of binders, we only used flycodes for binder ranking (relative quantification) in all three applications of this manuscript.

**Figure S7:**
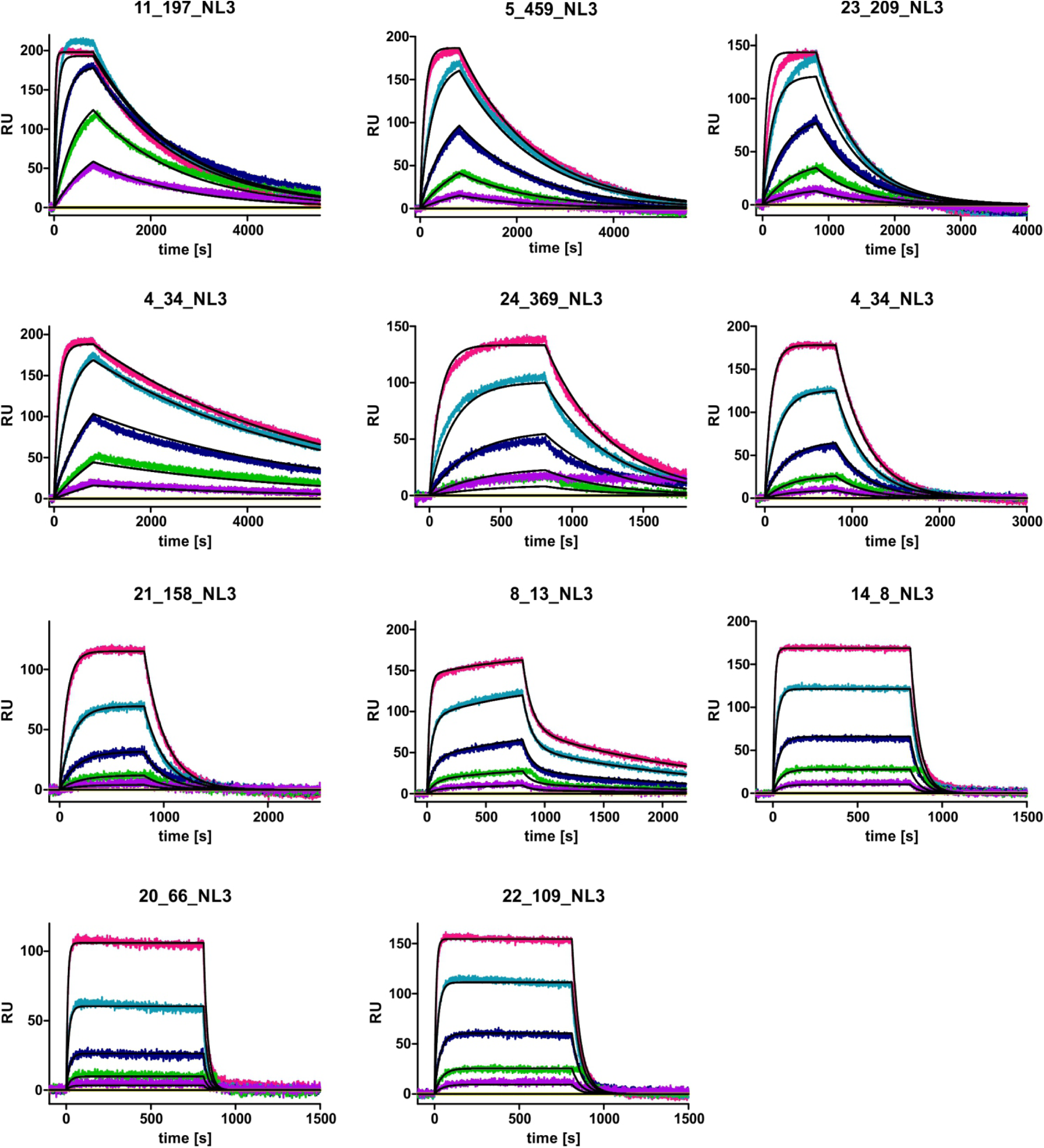
SPR analysis of MBP sybodies identified by NestLink. Data were measured with the ProteOn™ XPR36 Protein Interaction Array System (Bio-Rad) using biotinylated MBP (immobilized) and 81 nM, 27 nM, 9 nM, 3 nM, 1 nM of the purified sybodies.

**Figure S8:**
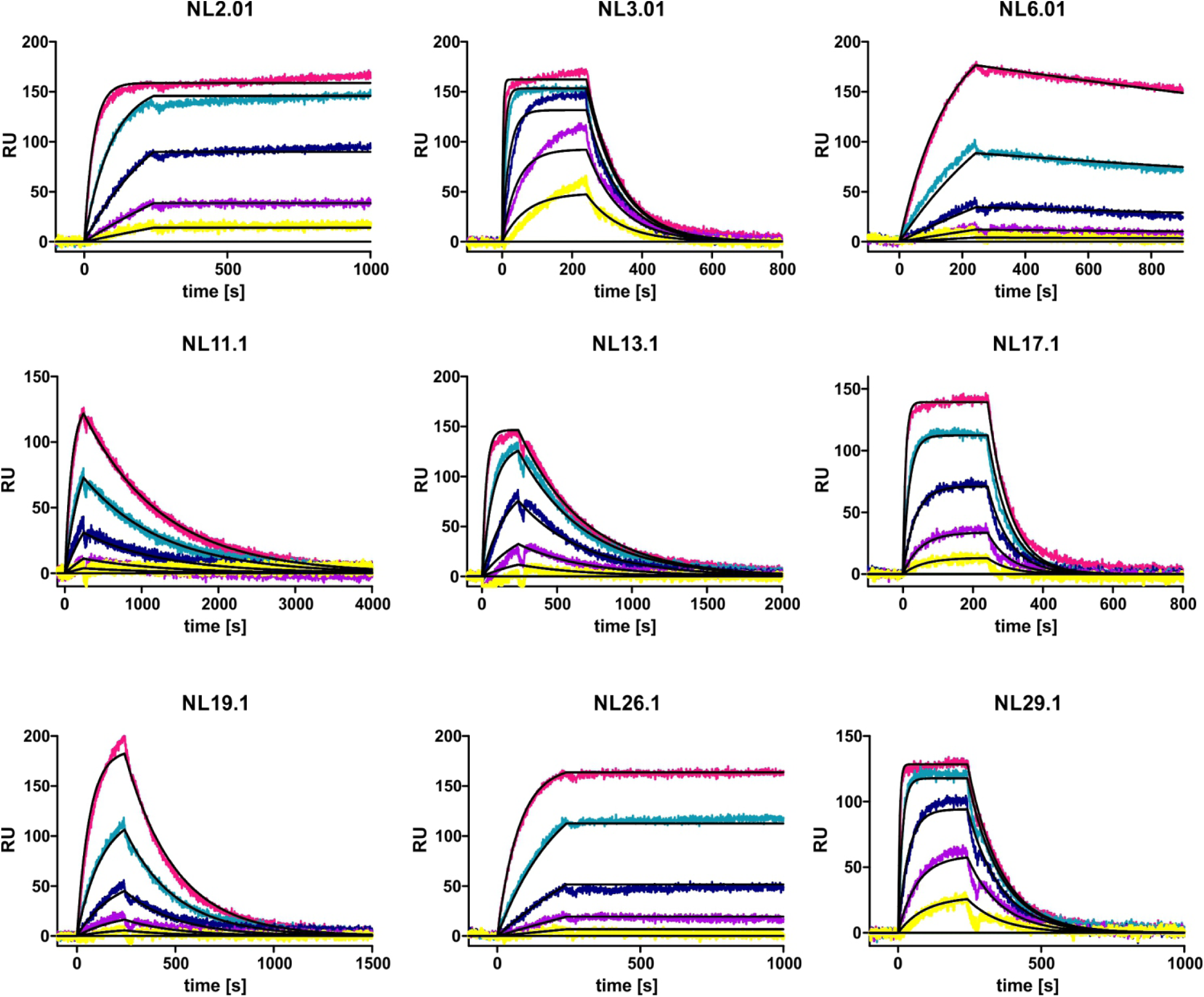
SPR analysis of nanobodies selected in solution against TM287/288. Data were measured with the ProteOn™ XPR36 Protein Interaction Array System (Bio-Rad) using biotinylated TM287/288 (immobilized) and the following concentrations of the purified nanobodies: 27 nM, 9 nM, 3 nM, 1 nM, 0.33 nM for NL2.01, NL13.1; 81 nM, 27 nM, 9 nM, 3 nM, 1 nM for NL3.01, NL6.01, NL11.1, NL19.1, NL26.1, NL29.1; 243 nM, 81 nM, 27 nM, 9 nM, 3 nM for NL17.1.

**Figure S9:**
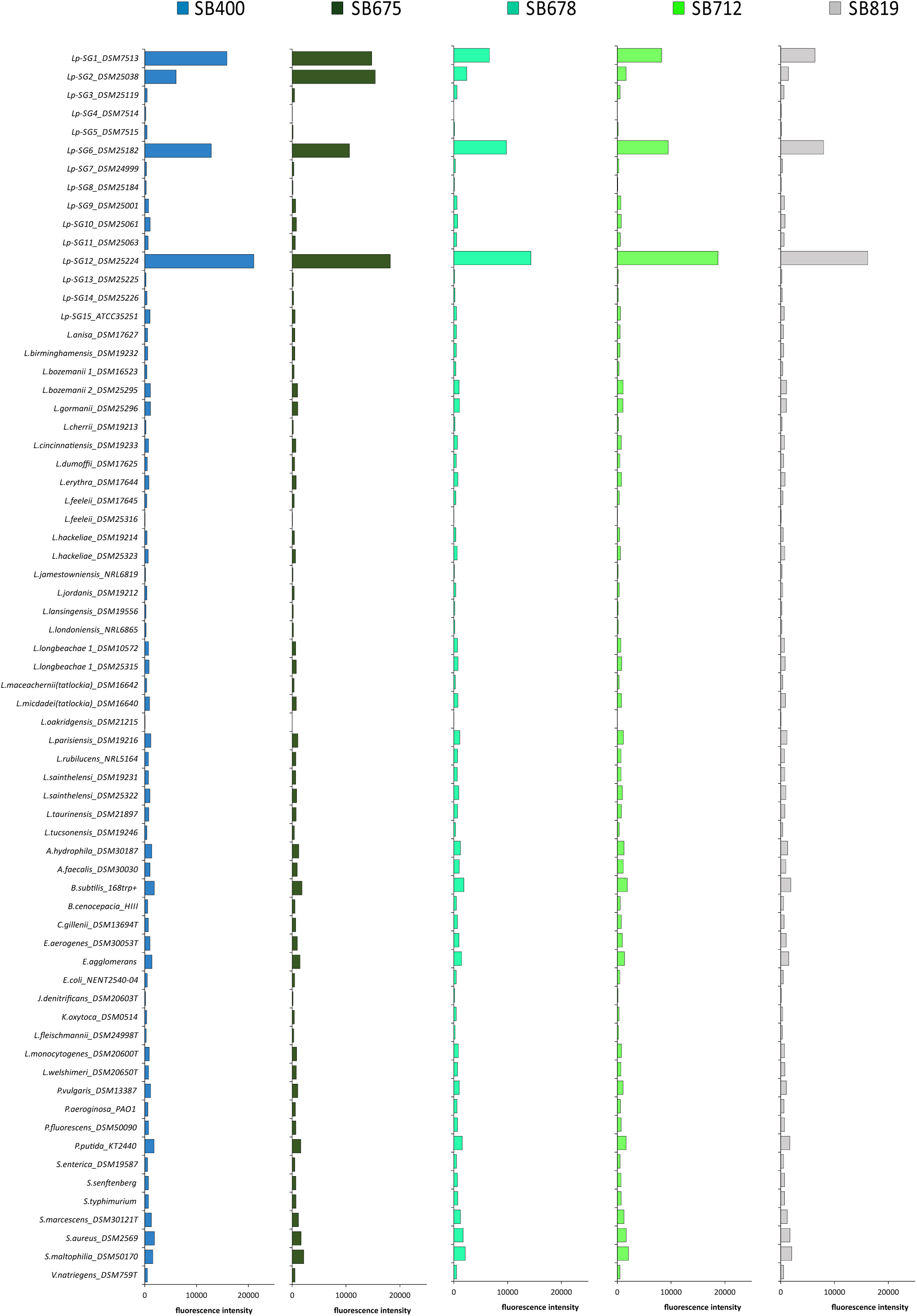
Flow cytometry screen. Detection of cell-surface binding on various bacterial strains for the sybodies recognizing MOMP of *Lp-SG6* (extended analysis of Fig. 6d).

**Figure S10:**
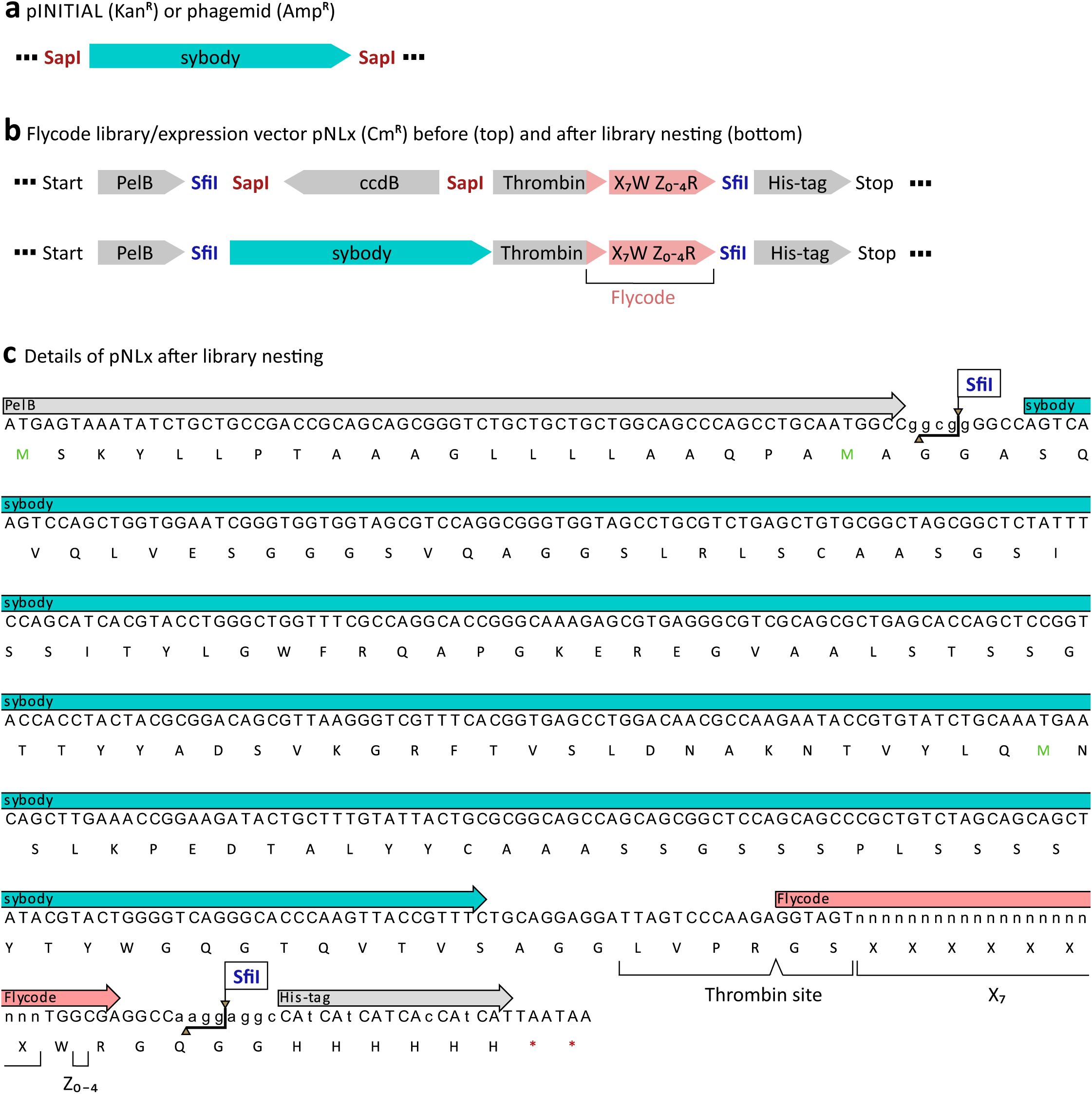
Vectors for library nesting. (**a**) Library members of interest (here sybodies) are cloned into an FX cloning initial vector^20^, from which they are excised using type IIS restriction enzyme SapI. (**b**) The flycode library is harbored on the *E. coli* expression vector pNLx. The library of interest is inserted via exchange of *ccdB* using SapI. (**c**) Open reading frame of pNLx after library nesting shown at the example of a sybody linked to an 11 amino acid flycode. SfiI cleavage is used to attach Illumina adaptors.

